# Radiation-reprogrammed glioma stem cells generate vascular-like cells to build a trophic niche driving tumor recurrence

**DOI:** 10.1101/2021.06.03.447005

**Authors:** Sree Deepthi Muthukrishnan, Riki Kawaguchi, Pooja Nair, Rachna Prasad, Yue Qin, Maverick Johnson, Nathan VanderVeer-Harris, Michael C. Condro, Alvaro G. Alvarado, Amy Pham, Raymond Gau, Qing Wang, Maria G. Castro, Pedro R. Lowenstein, Arjun Deb, Jason D. Hinman, Frank Pajonk, Terry C. Burns, Steven A. Goldman, Daniel H. Geschwind, Harley I. Kornblum

## Abstract

Treatment-refractory glioma stem and tumor cells exhibit phenotypic plasticity driving recurrence, but the underlying molecular mechanisms remain to be elucidated. Here, we employed single-cell and whole transcriptomic analyses to discover that radiation induces a dynamic shift in functional states of glioma cells allowing for acquisition of vascular endothelial-like and pericyte-like cell phenotypes. These vascular-like cells provide a trophic niche to promote proliferation of irradiated glioma cells, and their selective depletion results in reduced tumor growth post-treatment *in vivo*. Mechanistically, the acquisition of vascular-like phenotype is driven by increased chromatin accessibility and H3K27 acetylation in specific vascular gene regions post-treatment. Blocking P300 histone acetyltransferase activity reverses the epigenetic changes induced by radiation, and inhibits the phenotypic transition and tumor growth. Our findings highlight an important role for P300 histone acetyltransferase in treatment-induced plasticity and opens a new therapeutic avenue for preventing glioma recurrence.

**Significance:** Our study demonstrates that radiation therapy promotes glioma resistance by inducing vascular-like phenotypes in GSC that, in turn, aid in proliferation of the remaining tumor cells. This phenotype switch is mediated by P300 HAT, and inhibition of this enzyme is a potential therapeutic target for preventing glioma recurrence following radiation.

## Introduction

Glioblastoma (GBM) is a universally recurrent and lethal primary brain tumor that is highly resistant to standard chemo- and radiation therapy (1,2). Therapeutic failure and tumor relapse is partially attributed to pre-existing resistant fraction of glioma stem-like cells (GSC) that exhibit a high degree of plasticity and heterogeneity (3). Paradoxically, several studies have reported that chemotherapy with temozolomide (TMZ)- and radiation therapy-induced stress can themselves lead to dedifferentiation of tumor cells to a glioma stem cell-like (GSC) state, and these cells are termed induced-GSC (iGSC) and have been shown to promote tumor growth in xenograft models (4–7). A growing body of evidence also indicates that GSC switch from a Proneural to Mesenchymal state in response to ionizing radiation driving tumor invasion and resistance, and that they also transdifferentiate into vascular endothelial-like cells contributing to vascularization and tumor growth (8–14). GSC can also give rise to vascular pericytes that are essential for tumor growth and maintenance of blood tumor barrier (15). However, it is not clear whether therapy can induce the pericyte-like phenotype and whether they contribute to therapeutic resistance and relapse. While there has been tremendous progress on illuminating the mechanisms underlying GSC plasticity and heterogeneity in primary tumors, the precise molecular and epigenetic factors driving therapy-induced plasticity in treatment-refractory GSC and tumor cells that drive glioma recurrence remains to be elucidated.

In this study, we utilized single-cell and whole transcriptomic approaches to determine the contribution of radiation therapy in reprogramming glioma cells to adopt diverse phenotypic cell states. We found that clusters predominantly expressing vascular endothelial and pericyte/mesenchymal markers increase in size post-radiation, and that these clusters are enriched for gene sets associated with angiogenesis and vascular development. Using patient-derived gliomasphere cultures, orthotopic xenograft and murine GBM models, we demonstrate that radiation promotes the phenotypic transition of GSC to vascular endothelial- and pericyte like-cells, which, in turn influence tumor growth and recurrence post-treatment. Importantly, we uncovered that radiation alters chromatin accessibility in specific vascular gene regions resulting in their increased expression. The histone acetyltransferase (HAT) P300 mediates the acquisition of these vascular-like cell phenotypes in GSC and tumor cells in response to radiation and contributes to tumor relapse, opening up a new avenue for therapeutic targeting

## Results

### Single-cell transcriptomic sequencing reveals a dynamic shift in cellular states of glioma cells post-radiation

To determine the contribution of radiation to phenotypic plasticity in GBM, we performed single cell RNA-sequencing of a primary gliomasphere line exposed to a single dose of radiation at 8 Gy for 2- and 7-days. Integrated analysis of all the samples revealed 14 distinct clusters, a majority of which showed a dynamic shift in size (clusters 1, 2, 3, 5, 8, 10, 11), while a few increased (clusters 0, 4, 9) and some sharply reduced (clusters 6, 7, 12, 13) between 2- and 7-days post-radiation (Fig 1A-B). We next performed an unbiased analysis using co-expressed gene networks and clustering by Louvain community detection to annotate clusters based on functional cell states. 29 gene-signature modules were identified that were differentially expressed between the 14 clusters, and clustering divided them into two major fractions that further separated into 4 sub-groups (Fig 1C). Gene ontology (GO) analysis of the modules showed that subgroup S2 that contained clusters 6 and 7 (diminished post-radiation) were enriched for mitosis and DNA-repair processes. Subgroup S4, which comprised of clusters 0 and 4 (increased post-radiation) were highly enriched for embryonic development, vasculogenesis and mesenchymal stem cell differentiation modules. Subgroups S1 and S3 that contained clusters which either increased (9), reduced (12,13,14) or dynamically changed (3,10,11) post-radiation showed enrichment for mitosis, stress-response, homeostasis and embryonic development-related processes, indicating the diverse functional states of these cells (Fig 1C).

**Figure 1:**
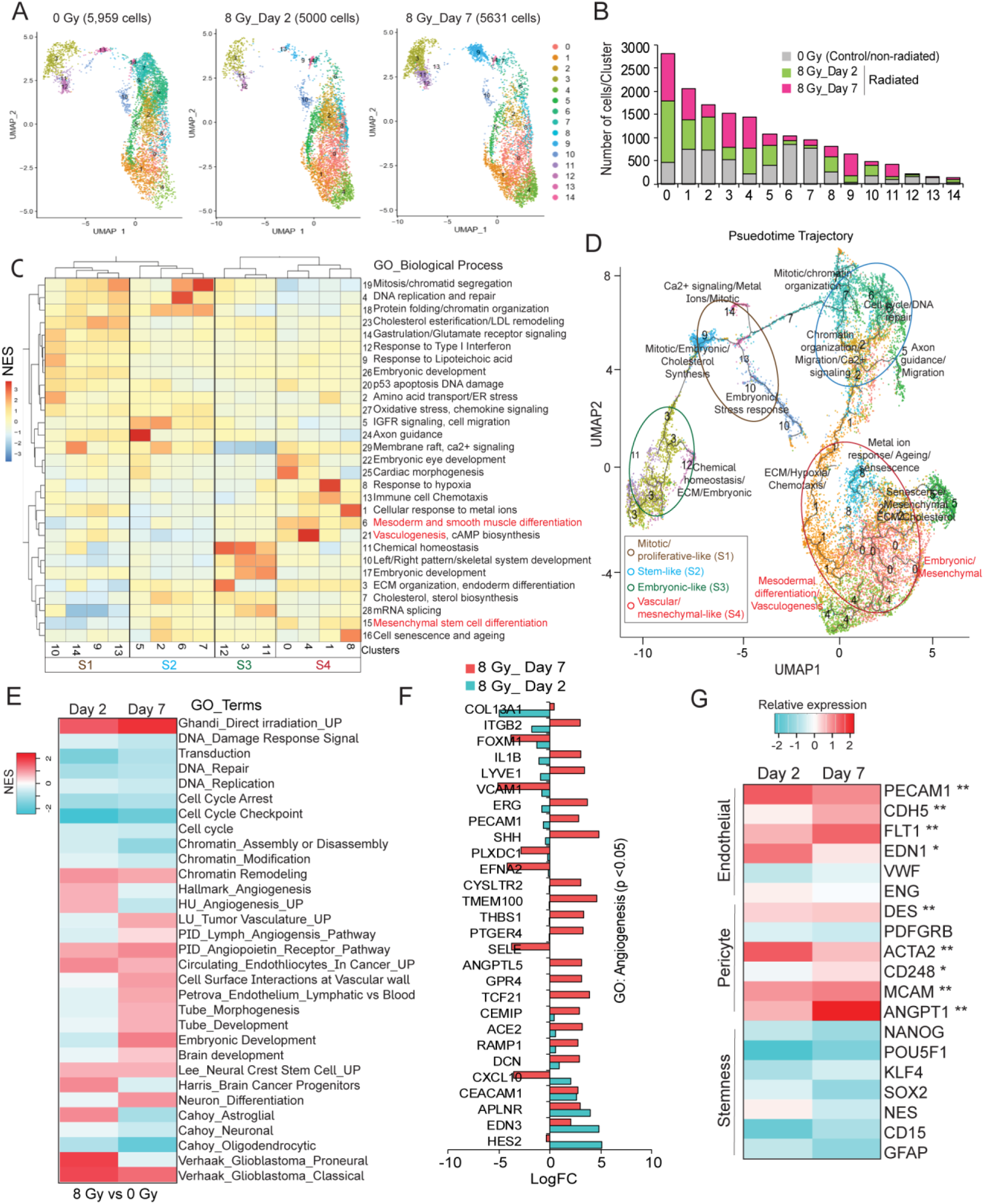
Single-cell sequencing reveals diverse functional states of radiated glioma cells. (A) UMAP plots show clustering of glioma cells from control (0 Gy) and 2-days and 7-days radiated (8 Gy) fractions. (B) Histogram shows the number of cells per cluster in each group. (C) Heatmap shows co-expressed gene network modules, Louvain clustering and EnrichR GO analysis of each cluster. Each subgroup based on clustering is highlighted by a black box. (D) UMAP plot of pseudotime trajectory analysis. Subgroups are highlighted in colored circles and their defined functional cell states are indicated in the inset. (E) Heatmap shows normalized enrichment score (NES) of gene sets in glioma cells 2- and 7-days post-radiation. (F) Graph shows LogFc expression of genes in the GO: Angiogenesis category in radiated vs control glioma cells. (G) Heatmap shows relative expression of endothelial, pericyte and stemness genes in control and radiated glioma cells. N=3, *, ** indicates p<0.05, and p<0.005, two-tailed t-test. See also Figure S1.

Pseudotime trajectory analysis with cluster 0 chosen *a priori* as an arbitrary starting point, and assigning highly enriched GO_terms to their respective clusters revealed the hierarchical sequence of these functional states. We found that on the one hand, subgroup S2 representing the stem cell-like state (in blue) branched into subgroups S1 (mitotic/proliferative-like state, in brown) and S3 (embryonic-like, in green), and on the other, gave rise to subgroup S4 (in red) enriched for vascular/mesenchymal-like processes (Fig 1D). To further validate these functional states, we examined for known markers of stem (glioma stem cell/GSC), vascular (endothelial and mesodermal), and mesenchymal (pericyte/neural crest) cells. We included the neural crest and mesodermal markers because pericytes, which have been shown to arise from GSC in tumors, are embryonically derived from neural crest/mesenchymal stem cells (NC-MSC) in the brain and vascular endothelial cells are derived from mesodermal progenitor cells (15–17). Consistent with the GO analysis results, cluster 0 from subgroup S4 highly expressed markers of GSC, NC-MSC and mature pericyte markers indicating that these may indeed represent mesenchymal/pericyte-like cell population. Interestingly, cluster 4 belonging to the same subgroup S4 showed increased expression of some mesodermal, and many mature endothelial markers suggesting that these maybe endothelial-like cells (**Fig S1A**). All clusters from subgroup S2 showed high expression of several GSC markers confirming that these are stem-like cells. Clusters (9,13,14) in subgroup S1 are enriched for mitotic markers and showed virtually no expression of GSC markers indicative of proliferating tumor cells. Clusters in subgroup S3 showed variable expression of GSC and embryonic markers suggesting that that these might be a transient cell population as they change dynamically in size between 2- and 7-day post-radiation (**Fig S1A**). These data indicate that a fraction of tumor cells and GSC that survive radiation-induced damage acquire the vascular endothelial and mesenchymal, pericyte-like cell states.

To examine if the *in vitro* cell states are replicated *in vivo*, we also performed scRNA-sequencing on tumor cells isolated from radiated and control xenografts generated in NSG mice. Integrated analysis of the samples identified 15 clusters in total, of which clusters 1,2-4,9,10, 12 and 13 increased in frequency, and clusters 0, 5, 6,7 and 11 diminished in radiated tumors. **(Fig S1B-C**). Similar to *in vitro* findings, Louvain community detection clustering revealed 2 major fractions and 4 subgroups (**Fig S1D**). However, subgroup 3 was further divided into two: S3a and S3b as they showed enrichment for different sets of modules. Overall, clusters with greater frequency in radiated tumors were enriched in adhesion/junction assembly, cGMP signaling, angiogenesis, epithelial-mesenchymal transition, and migration-related processes, whereas clusters that diminished in size expressed modules enriched in embryonic eye development, ion homeostasis, migration, and stem cell differentiation (**Fig S1D**). Pseudotime trajectory and known marker analysis revealed that clusters (4,9) from subgroup S1 (blue) had highest expression of GSC markers representing the stem-like cells. Subgroup S3a (red) with clusters (10,13) showed increased expression of mature endothelial markers, and S3b with clusters (2,3,5) represented the mitotic and proliferating tumor cells. Subgroup S4 (red) containing cluster 1 displayed high levels of NC-MSC and pericyte markers, and all these clusters increased in size post-radiation (**Fig S1E, F).** Together, these data indicate that refractory tumor cells acquire stem-like, endothelial-like and mesenchymal/pericyte -like phenotypes in tumor xenografts, essentially replicating the *in vitro* findings.

### Whole-transcriptomic sequencing confirms the enrichment of angiogenic and vascular markers in gliomaspheres post-radiation

Next, we sought to determine if radiation-induced functional states observed in single cell studies are discernible in patient-derived gliomasphere cultures by bulk RNA-sequencing. Hierarchical clustering of normalized gene expression showed that radiated cells at day 2 and day 7 clustered separately from each other, and from control cells **(Fig S1G, H).** As expected, gene set enrichment analysis (GSEA) showed enrichment of gene ontology (GO) terms related to radiation, and reduction in gene sets associated with cell cycle and DNA repair in both day 2 and day 7 radiated cells (Figure 1E). Furthermore, in line with single cell studies, we found enrichment of gene sets related to angiogenesis, embryonic development, NC/MSC and mesenchymal transition, and downregulation of proneural and neuronal gene sets in radiated gliomaspheres. Closer examination of the genes associated with these processes showed significant enrichment of angiogenesis genes (*p<0.05, Benjamini-Hochberg adjusted*), and a small subset of mesenchymal genes from the Verhaak_Glioblastoma_Mesenchymal geneset in 7-day radiated gliomaspheres (Fig 1F**, Fig S1I**). Quantitative RT-PCR and immunostaining confirmed the enrichment in vascular markers that included mature endothelial (PECAM1, CDH5, FLT1, EDN1) and pericyte (DES, ACTA2, ANGPT1, MCAM, CD248) genes in day 7 radiated gliomaspheres (Fig 1G**, Fig S1J**). These markers were not significantly upregulated in day 2 gliomaspheres, corroborating the findings from bulk RNA sequencing. This data suggests that radiation promotes vascular gene expression in gliomaspheres.

### Radiation-resistant glioma cells exhibit increased endothelial and pericyte marker expression

Standard radiation therapy for GBM patients involves a fractionated dose regimen of 2 Gy for 30 days (18). However, our transcriptomic analyses were performed with a single high dose of radiation, 8 Gy. Therefore, we assessed whether fractionated radiation (2 Gy, x4) also induces vascular-like phenotypes in gliomaspheres. Quantitative RT-PCR analysis of both fractionated and single dose radiated gliomaspheres showed significant increase in endothelial and pericyte markers (Fig 2A). We also found that the increase in vascular marker expression by radiation was not dose-dependent, and that both low (2 and 4 Gy) and high (8 and 10 Gy) dose were effective at inducing the expression of the majority of the vascular genes assessed (**Fig S2A**). GSC and mesenchymal markers were not significantly upregulated by fractionated and single-dose radiation **(Fig S2B)**.

**Figure 2:**
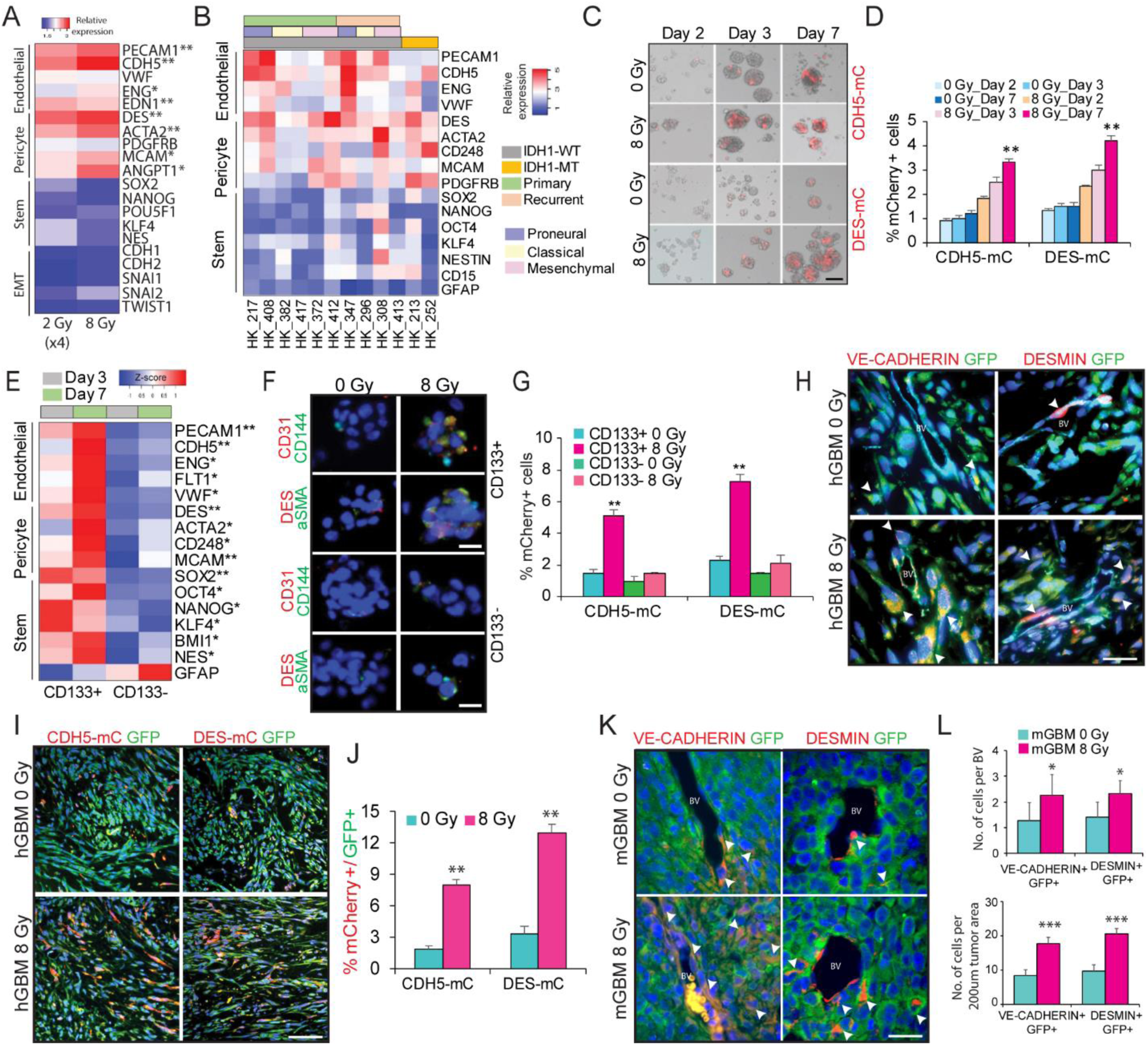
Radiation promotes endothelial and pericyte marker expression in glioma cells. (A) Heatmap shows relative expression of endothelial, pericyte, stemness and EMT genes in control (0 Gy), fractionated (2 Gy X4) and single dose (8 Gy) radiated glioma cells. (B) Heatmap shows relative expression of endothelial, pericyte and stemness markers in control and radiated glioma cells from multiple patient-derived gliomasphere lines. (C, D) Representative images of mCherry expression in control and radiated cells. Graph shows flow-cytometric quantitation of percentage of mCherry+ cells in each condition. (E) Heatmap shows relative expression of endothelial, pericyte and stemness genes in CD133+ (GSC) and CD133- (non-GSC) fractions 3- and 7-days post-radiation. (F) Immunostaining images of endothelial (CD31, CD144/VE-CADHERIN) and pericyte (DES, aSMA) markers in control and radiated CD133+ (and CD133-fractions of glioma cells. Scale bars, 50μm. (G) Flow cytometric quantitation of percentage of mCherry+ cells in control and radiated CD133+ and CD133-fractions. (H) Immunostaining images of VE-CADHERIN and DESMIN markers in GFP+ tumor cells in control and radiated tumor xenografts. Scale bars, 50μm. White arrows point to GFP+ marker+ cells in each image. (I,J) Immunostaining images of mCherry and GFP in control and radiated tumor xenografts. Graph shows flow-cytometric quantitation of percentage of mCherry+ cells normalized to total number of GFP+ tumor cells in each group. N=2, and ** indicates p<0.005 derived from Welch’s t-test. Scale bars, 100μm (K, L) Immunostaining images of VE-CADHERIN and DESMIN in GFP+ tumor cells in mouse GBM model. Arrows point to marker+ GFP+ cells in each group. Scale bars, 50μm. Graphs show quantitation of number of GFP+ marker+ cells in blood vessel (BV) and tumor mass. N=3, * and *** indicates p<0.05 and p<0.0005 derived from Welch’s t-test. Error bars represent s.d. of the mean in (D, G), N=3, * and ** indicates p<0.05 and p<0.005, two-tailed t-test. See also Figure S2.

To determine the generalizability of our findings, we used the single high dose 8 Gy to verify if radiation promotes vascular marker expression in multiple patient-derived gliomasphere lines that included the 3 major molecular subtypes: Proneural, Mesenchymal and Classical (19), and both primary and recurrent IDH1-wild type and mutant GBM. We found significant upregulation of vascular markers, albeit to a varying degree, in all gliomasphere lines examined irrespective of the molecular subtype and mutational status. Interestingly, Proneural lines (HK_217, HK_408, HK_347) showed the highest expression for endothelial markers (PECAM1, CDH5), and the Mesenchymal lines (HK_372, HK_412, HK_308) exhibited the highest pericyte marker expression (DES, ACTA2). Although the primary GBM lines did not show an increase in GSC markers, some of the recurrent GBM lines showed an increase in GSC markers including SOX2, NANOG and OCT4 post-radiation (Fig 2B). These findings indicate that radiation can promote stem-like and vascular-like marker expression, but the acquisition of stem-like state may be restricted to a subset of gliomas.

To quantify the extent of radiation-induced phenotype switching, we generated lentivirus constructs containing endothelial (VE-CADHERIN/CDH5) or pericyte (DESMIN) promoter to drive expression of fluorescent mCherry reporter. We found that maximal reporter activation from both promoters occurs 3- and 7-days post-radiation (Fig 2C-D). Flow cytometric analysis using previously validated endothelial (CD31, CD144) and pericyte markers (CD146, CD248) also showed increased percentage of double-positive cells 3- and 7-days post-radiation (**Fig S2C).** Co-expression of both reporters showed that the majority of cells did not display overlapping expression of VE-CADHERIN-mCherry and DESMIN-GFP in both control and radiated gliomaspheres (**Fig S2D**). Immunostaining and qRT-PCR analysis of FACS sorted reporter-positive and negative fractions also indicated that endothelial and pericyte markers are expressed by distinct cells in gliomaspheres supporting the results from single-cell studies that they constitute different cell states/phenotypes **(Fig S2E-F).**

In order to distinguish between the possibilities that radiation: a) induces the vascular-like phenotypes versus b) promotes expansion of pre-existing vascular-like cells, we preemptively depleted the reporter positive cells by FACS. Radiation of reporter-negative fractions activated both VE-CADHERIN and DESMIN reporters, but no activation was seen in non-radiated gliomaspheres (**Fig S2G-H**). We also extended these findings by examining reporter activation in multiple patient-derived gliomasphere lines. Consistent with the gene expression data, while gliomaspheres from the Proneural (PN) and Classical (CL) subtypes showed similar expression levels for both reporters, the Mesenchymal (MES) subtype showed significantly higher pericyte reporter expression compared to endothelial reporter (**Fig S4I**), indicating that glioma cells from specific molecular subtypes may exhibit differential propensity for acquiring endothelial-like and pericyte-like states in response to radiation.

### GSC exhibit higher propensity for vascular-like phenotype switching than non-GSC in response to radiation

Prior studies have demonstrated that CD133+ GSC transdifferentiate into endothelial cells and pericytes (12,15,20). To determine if radiation promotes vascular-like phenotype switching of GSC or non-GSC tumor cells, we also utilized CD133 (PROM1) as the GSC marker to separate the stem-like fraction from non-stem like cells (21). Because this GSC marker is not informative in all tumors and cultures, we validated its significance in the cell line being used, HK_408 with limiting diluting assay and orthotopic transplantation of a small number (1000 cells) of CD133+ and CD133-cells into mice (**Fig S2J-K).** Radiation of either the CD133+ or CD133-fractions showed that significantly increased endothelial and pericyte marker expression was restricted primarily to the CD133+ GSC fraction at 3 and 7-days post-radiation. GSC markers were also highly upregulated in the CD133+ GSC fraction relative to CD133-tumor cells (Fig 2E). Immunostaining of endothelial (CD31/VE-CADHERIN) cells and pericyte (DESMIN/aSMA) markers, and reporter expression confirmed that GSC exhibit greater capacity for vascular-like phenotype switching than non-GSC post-radiation (Fig 2F-G). These findings indicate that at least for this gliomasphere line, that radiation-induced vascular-like phenotype is isolated to the stem cell fraction.

### Radiation promotes vascular-like phenotypes in orthotopic xenograft and murine GBM models

To determine if radiation promotes vascular-like phenotype switching in glioma cells *in vivo*, we first generated orthotopic xenograft tumors using the gliomasphere line HK_408 infected with a Firefly-Luciferase-GFP lentivirus. Tumor-bearing mice were exposed to a single dose of 8 Gy radiation. Immunostaining for vascular markers showed an increase in tumor cells co-expressing GFP with VE-CADHERIN or CD31 (endothelial) and DESMIN or αSMA (pericyte) predominantly in the tumor mass, and only a very rare number of cells in the vessels, in radiated tumors compared to non-radiated tumors (Fig 2H **and S2L**). To quantitatively measure *in vivo* vascular-like phenotype switching, we generated xenografts with tumor cells transduced with VE-CADHERIN (CDH5-mC) and DESMIN (DES-mC) reporters. Tumor-bearing brains from control and radiated groups were harvested 2 weeks after radiation, and analyzed by immunostaining and flow cytometry for the presence of GFP+ mCherry+ cells (Fig 2I). Immunostaining for GFP (tumor) and mCherry (vascular-like cells) revealed an increase in co-labelled cells and flow cytometric quantitation showed that this increase was significant for endothelial- (8% vs 2% control, * p<0.005) and pericyte- (13% vs 3% control, **p<0.005) like cells in radiated tumors (Fig 2J). These findings, consistent with *in vitro* studies, indicate that radiation promotes the acquisition of endothelial-like and pericyte-like phenotypes in glioma cells *in vivo*.

Next, we asked whether the presence of an intact immune system affects the acquisition of vascular-like phenotypes in tumor cells post-radiation in a syngeneic mGBM model (22). We first verified that mGBM cultures showed enrichment of vascular genes post-radiation (**Fig S2M)**. Examination of tumors showed that there was co-staining of GFP (tumor cells) with endothelial (VE-CADHERIN) and pericyte (DES) markers post-radiation. Quantification of GFP+ marker+ cells revealed that there was significant increase in vascular-like cells in radiated mice, most predominantly in the tumor mass, but only a small yet significant increase in vessels (Fig 2K, L **and Fig S2N).** This indicates that radiation-induced vascular-like phenotype switching is not inhibited by an intact immune microenvironment as would exist in human tumors.

### Radiation-induced glioma endothelial cells (iGEC) display phenotypic characteristics of normal vascular endothelial cells

Typically, the identity and behavior of normal vascular endothelial cells (VEC) is ascertained by their ability to uptake labelled low-density lipoproteins (LDL) and form tubular networks on GFR-Matrigel mimicking the blood vessels (23–25). Earlier studies have shown that glioma-derived endothelial cells possess these characteristics (12,26,27). We therefore asked if radiation-induced glioma endothelial cells, referred to henceforth as “*iGEC*” also exhibit characteristics of normal VEC. We used human umbilical vein endothelial cells (HUVEC) as a reference for normal VEC to assess these phenotypes. Similar to HUVEC, radiated glioma cells also showed tubular-network formation on Matrigel, and increased uptake of DiI-Ac-LDL compared to non-radiated cells (Fig3A-D**, Fig S3A-B**).

**Figure 3:**
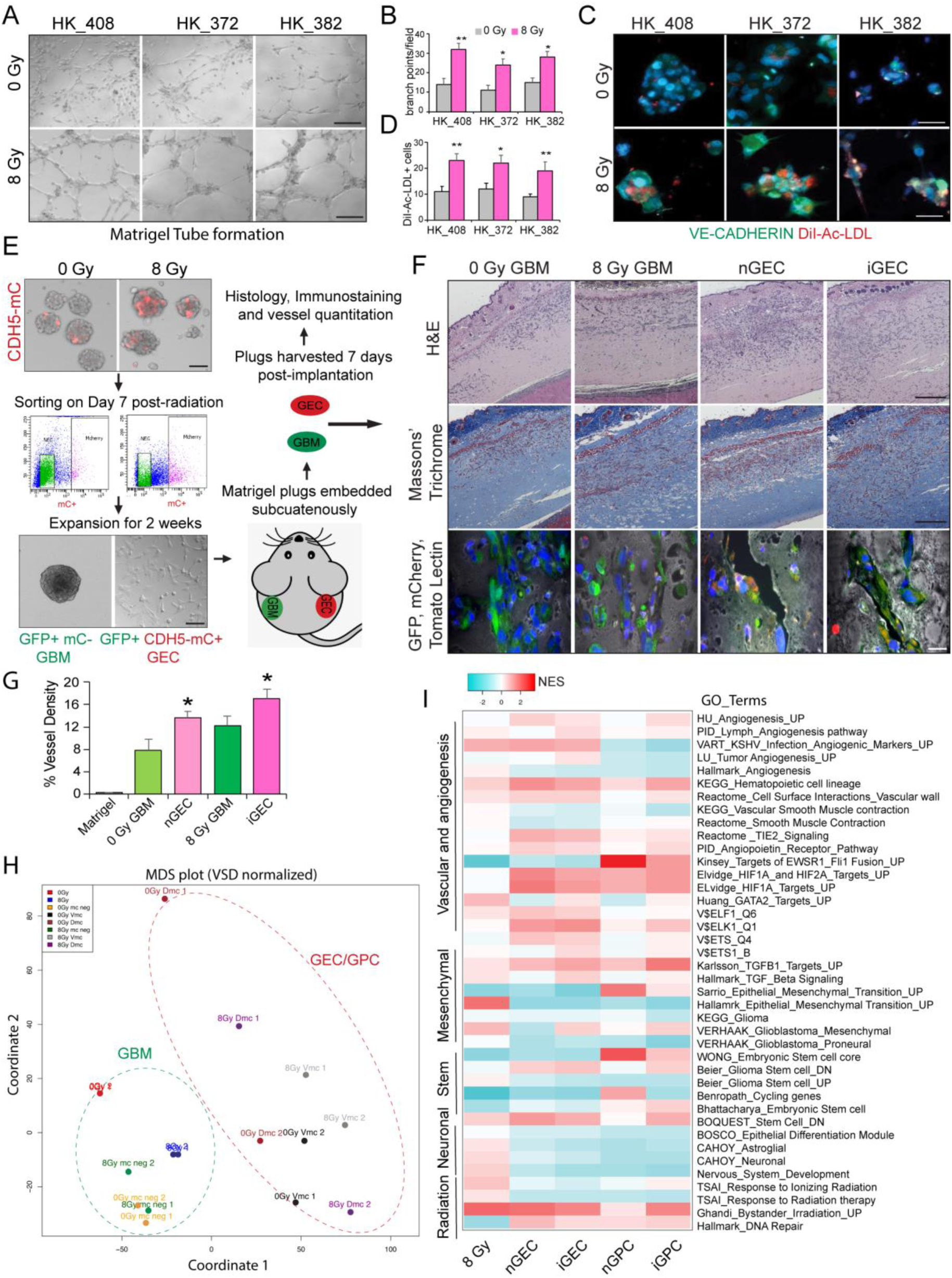
iGEC display typical characteristics of normal vascular endothelial cells. (A-D) Tubular network formation on matrigel and immunostaining of VE-CADHERIN and Di-Ac-LDL in control and radiated glioma cells. Scale bars, 100μm. Graphs show quantitation of branch points and number of Di-Ac-LDL+ cells per field in each group. Error bars represent mean±SD, and * and ** indicates p<0.05 and p<0.005, Welch’s t-test. (E) Schematic outlines the matrigel plug in vivo angiogenesis assay. (F) Images show H&E and Masson’s trichrome staining. Scale bars, 500μm. Representative images of immunostaining of mCherry (GEC, red), GFP (tumor cells, green) and Tomato Lectin (vessels, grey). Scale bars, 20μm. (G) Graph shows quantitation of vessel density in each group. Error bars represent mean ± SD and *, ** indicates p<0.05 and p<0.005, two-tailed Students t-test. (H) MDS plot shows clustering of tumor and transdifferentiated cells. (I) Heatmap shows NES of gene sets in each group. See also Figure S3.

We next performed Matrigel plug *in vivo* angiogenesis assay using FACS sorted reporter-negative tumor cells (mCherry-) and or reporter-positive iGEC or nGEC (CDH5-mCherry+) to determine if they integrate into host vasculature or induce angiogenesis when embedded subcutaneously (Fig 3E). Histological examination with hematoxylin & eosin (H&E) and Masson’s Trichrome staining revealed vessel formation in both GBM and GEC-embedded plugs. Immunostaining revealed a few mCherry+ cells lining along the vessels labelled with Tomato Lectin only in iGEC and nGEC groups indicating that some vascular-like cells may integrate into host vasculature (Fig 3F). Quantitation of Trichrome stained sections also showed that both iGEC and nGEC groups had significantly higher vascular density than their corresponding tumor fractions (Fig 3G).

To validate that a) the angiogenic and vascular gene signatures are restricted to vascular-like cells, and b) they are molecularly distinct from the rest of the tumor cells, we performed RNA-sequencing on sorted reporter+ (iGEC/GPC and nGEC/GPC, mCherry+) and reporter- (mCherry-) tumor cells from control and radiated gliomaspheres. Multi-dimensional scaling (MDS) and hierarchical clustering of gene expression showed that tumor cells and vascular-like cells clustered separately (Fig 3H). Differential gene expression analysis also revealed significant number of up and down-regulated genes between tumor cells versus vascular-like cells (**Fig S3C**). Examination of canonical endothelial (CD31 and VE-CADHERIN) and pericyte (DES and ACTA2) markers confirmed the enrichment of these transcripts in GEC and GPC fractions, respectively (**Fig S3D**). GSEA also showed increase in vasculature and angiogenesis related gene sets in both non-induced and induced GEC and GPC fractions compared to their respective tumor fractions (Fig 3I). GPC fractions were also enriched for smooth muscle- and mesenchymal-gene signatures consistent with their mesenchymal identity (Fig 3I). We also observed downregulation of stem cell, neuronal, glial and nervous system development-related gene sets in the vascular-like cell fractions relative to tumor cells, further confirming that these vascular-like cells are molecularly distinct from their precursor tumor cells, and exhibit some vascular-like characteristics.

### iGEC and iGPC provide a trophic niche to support tumor growth post-radiation

Given that only a small number of iGEC and iGPC contribute to vessel formation, and majority are located in the tumor mass, we asked whether they provide a trophic niche to promote tumor growth following treatment. To test this hypothesis, we first collected conditioned media (CM) from the sorted and cultured non-induced (nGEC/nGPC) and induced (iGEC/iGPC) vascular-like cells (Fig 4A). Addition of CM from iGEC/iGPC to radiated tumor cells significantly promoted their proliferation, whereas it did not show significant growth promoting effect on non-radiated tumor cells (* p<0.05, N=4, one way-ANOVA). CM from unsorted GBM cells did not promote growth of radiated tumor cells (Fig 4B). We also performed this assay on control and radiated tumor cells isolated from tumor xenografts, and obtained similar results, confirming that the vascular-like cells provide trophic support to radiated tumor cells (**Fig S4A**). These findings indicate that iGEC and iGPC secrete factors that support the growth of radiated glioma cells.

**Figure 4:**
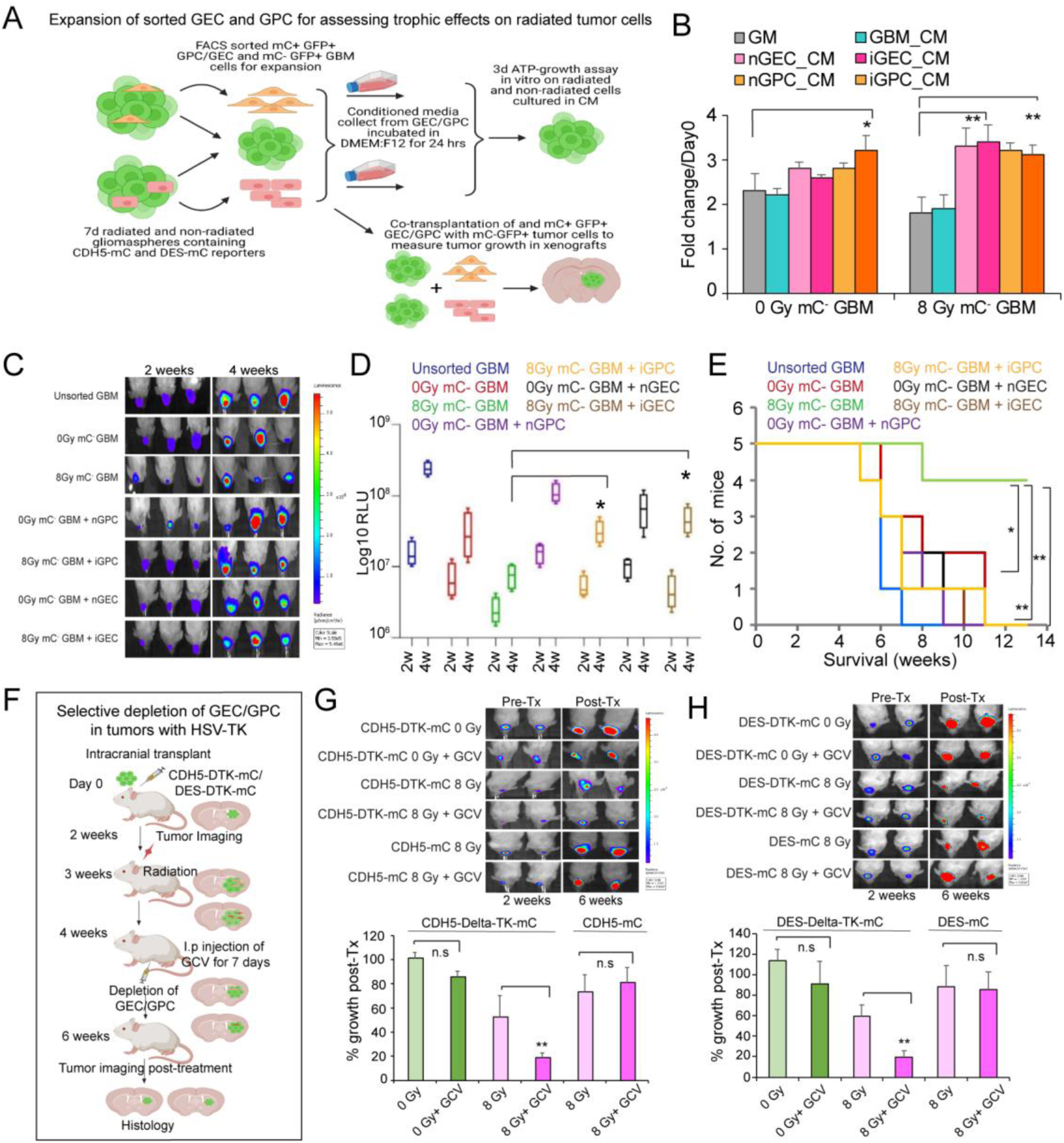
iGEC and iGPC provide trophic support to promote tumor growth post-treatment. (A) Schematic outlines the conditioned media (CM) and co-transplantation tumor growth experiments. (B) Graph shows quantitation of proliferation of control and radiated tumor cells in GEC and GPC conditioned media (CM). Error bars represent mean±SD, N=3, and *, ** indicates p<0.05 and p<0.005, two-tailed t-test. (C,D) Images of mice showing tumor growth at 2 and 4 weeks. Box plots show quantitation of tumor growth by luminescence in each group. N=5 mice per group and * indicates p<0.05, one-way ANOVA and post-hoc t-test. (E) Kaplan-Meier survival curve of mice transplanted with tumor cells and GEC/GPC. * and ** indicates p<0.05 and p<0.005, Log-rank test. (F) Schematic outlines the depletion of GEC/GPC using HSV-TK method in tumor xenografts (G,H) Images of mice showing tumor growth at 2 and 4 weeks. Box plots show the quantitation of tumor growth by luminescence. * indicates p-value <0.05 and n.s. not significant, one-way ANOVA and post-hoc t-test. See also Figure S4.

To assess the potential trophic function of iGEC and iGPC *in vivo*, we employed two complementary strategies. First, we performed co-transplantation of control and radiated reporter-negative GFP-Luciferase expressing tumor cells with their respective GFP-luciferase negative, mCherry-expressing GEC and GPC fractions that in a 1:1 ratio, and assessed tumor growth at 2 and 4 weeks after transplantation (Fig 4A). Remarkably, co-transplanted iGEC and iGPC significantly enhanced the growth of radiated tumor cells (reporter, mC^-^) compared to tumor cells transplanted alone, corroborating the *in vitro* findings (Fig 4C-D **and Fig S4B**). Survival analysis also showed that the co-transplanted mice exhibited signs of morbidity significantly earlier than mice transplanted with tumor cells alone (Fig 4E).

Next, we performed selective depletion of iGEC/iGPC *in vivo* using the HSV-TK-Ganciclovir cell ablation strategy by driving expression of HSV-TK under the control of either VE-CADHERIN (CDH5-HSVTK-mCherry) or DESMIN (DES-HSV-TK-mCherry) (Fig 4F). After verifying tumor formation, mice were radiated with a single dose of 8 Gy, and injected with GCV every day for one week to allow depletion of reporter+ cells expressing HSV-TK. Quantification of tumor growth pre- and post-radiation and GCV administration revealed that depletion of either iGEC and iGPC markedly reduced tumor growth after radiation (Fig 4G-H). On the other hand, depletion of nGEC and nGPC did not show a significant effect on non-radiated tumors. We also did not see growth inhibition of tumors transduced with reporter constructs lacking HSV-TK when administered with GCV indicating that only vascular-like cells expressing HSV-TK were selectively depleted resulting in growth inhibition **(Fig S4C, D).** We also utilized another cell ablation strategy by expressing Diphtheria Toxin Receptor (DTR) to selectively deplete vascular-like cells. Since human cells are known to express the DT receptor, HB-EGF, we first ensured that the protein is not expressed by our cell line, and also verified that Diphtheria toxin (DT) treatment selectively ablated the DTR-expressing cells (**Figure S4E-F**). Similar to HSV-TK depletion, mice-bearing tumors were radiated and administered with DT in 3 doses over the course of one week to ablate GEC/GPC **(Fig S4G)**, and found significant inhibition in tumor growth (**Fig S4H-J**). Collectively, the *in vitro* and *in vivo* findings strongly suggest that vascular-like cells provide a trophic niche to tumor cells and promote recurrence after radiation.

### Radiated gliomaspheres exhibit increased chromatin accessibility and H3K27Ac in specific vascular gene regions

Transition between cellular states requires epigenetic and transcriptional reprogramming driven by alterations in chromatin structure and accessibility (28,29). We therefore asked if radiation-induced vascular-like phenotype involves changes in chromatin accessibility, especially in vascular gene regions. To test this hypothesis, we performed ATAC-sequencing on control and radiated gliomaspheres 2-days post-radiation. We chose the 2-day time point to determine changes that occur in the surviving fraction of tumor cells immediately after radiation-induced damage and prior to significant induction of vascular marker expression. Although there was a trend towards reduced chromatin accessibility (as shown by overall peak counts) in radiated tumor cells, this difference was not significant (N=8 samples per condition and 3 independent experiments, p=0.3, two-tailed t-test (Fig 5A)). As shown in the graph, there was great variability between the peak numbers obtained from 3 independent experiments limiting any overarching conclusions regarding altered global chromatin accessibility in response to radiation. This was also reflected in percentage of peak distribution across various genomic regions, which did not show any overt differences (Fig 5B).

**Figure 5:**
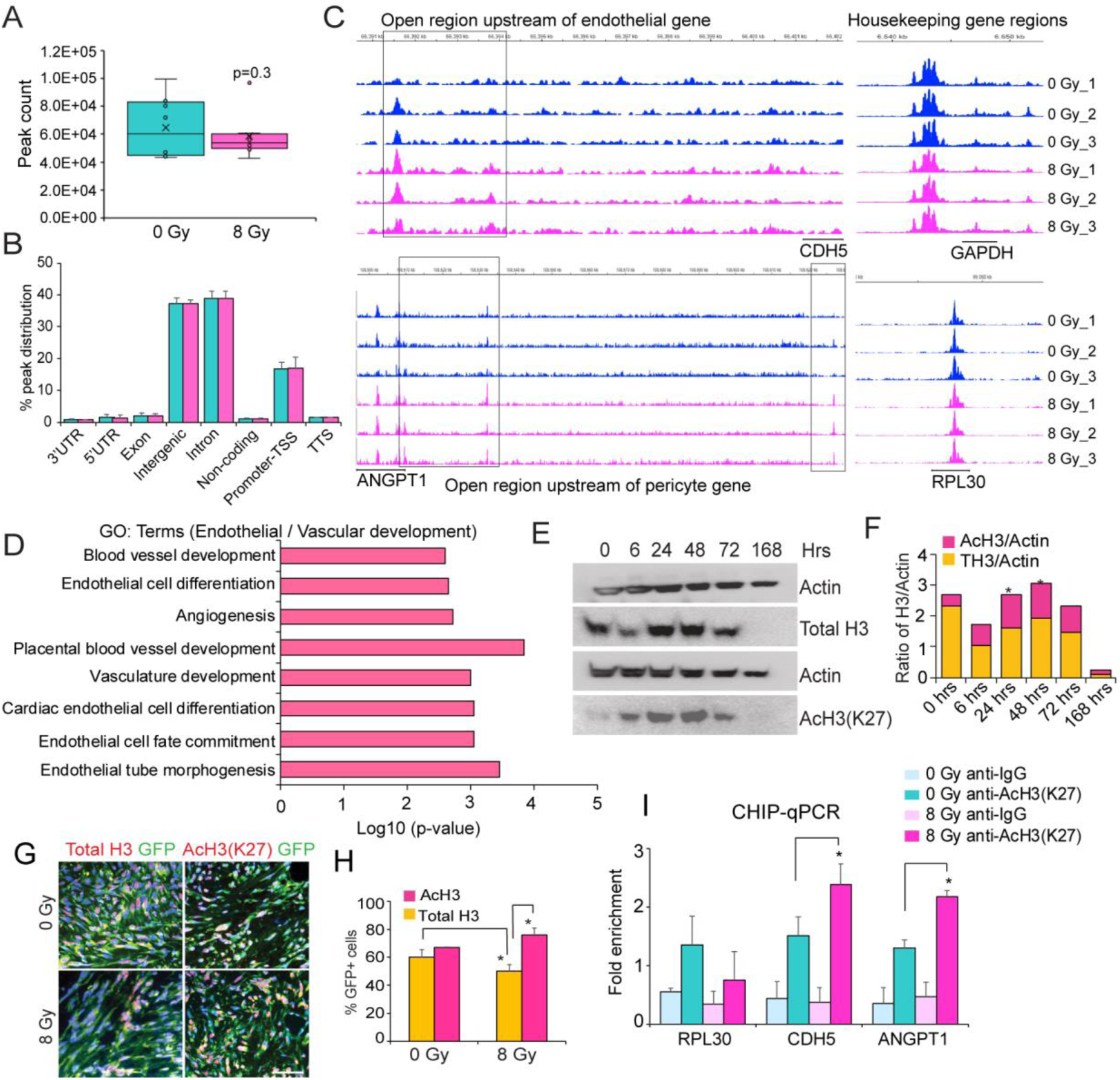
Radiation alters chromatin accessibility in vascular gene regions. (A,B) Total peak count and peak distribution across various genomic regions in 2-day control and radiated gliomaspheres. N=8 samples per group, and from 3 independent experiments. P=0.3 derived from two-tailed t-test. (C) Differentially open regions (highlighted in black boxes) upstream of vascular genes (CDH5, endothelial and ANGPT1, pericyte), and housekeeping genes (RPL30 and GAPDH) in non-radiated and radiated cells. (D) Gene ontology analysis shows enrichment of vascular and blood vessel development related gene terms in radiated gliomaspheres. (E, F) Immunoblot images of total and Ac-histone 3 (AcH3, Lysine K27) in control and radiated cells. Quantitation of protein levels normalized to Actin is shown in the graph, N=2 and * indicates p<0.05, two-tailed t-test. (G, H) Immunostaining images of Total H3 (red), AcH3 (red) and GFP (green) in control and radiated tumor xenografts. Quantitation of percentage of positive cells is shown in the graph, N=3 and * indicates p<0.05, two-tailed t-test. Scale bars, 100um. (I) Graph shows fold enrichment of vascular gene regions (CDH5 and ANGPT1) immunoprecipitated with anti-H327Ac and control anti-IgG antibody in control and radiated cells. RPL30 was used as a positive control. N=2 independent experiments and * indicates p<0.05 derived from one-way ANOVA. See also Figure S5.

Next, we examined if vascular genes that showed increased expression post-radiation displayed altered chromatin accessibility. Differential analysis revealed specific sites in vascular endothelial (VE-CADHERIN /CDH5) and pericyte (ANGPT1) genes to be more open (indicated by increase in peak size in 3 representative samples) in radiated glioma cells (Fig 5C). This difference in peak size was not reflected in all genes, and as an example, we show that the two housekeeping genes, RPL30 and GAPDH did not display altered accessibility between control and radiated gliomaspheres (Fig 5C). We also found that there are known H3K27Ac and transcription factor binding sites (TFBS) in these genomic locations in CDH5 and ANGPT1 genes in the UCSC genome browser, indicating that these regions may be important for their transcriptional activation (**Fig S5A**). Gene Ontology (GO) analysis of all the differentially open peaks revealed significant enrichment of terms associated with vascular development, endothelial differentiation, mesenchymal and stem cell development (Fig 5D and **Fig S5B**). HOMER motif analysis also revealed significant enrichment of vascular endothelial specification and mesenchymal transition-associated transcription factor motifs in radiated gliomaspheres (**Fig S5C).** These results suggest that radiation increases chromatin accessibility specifically in certain genomic regions of the vascular genes.

Histone acetylation is a key determinant of chromatin accessibility (30,31). We wondered whether radiation altered histone acetylation levels leading to changes in chromatin accessibility in vascular genes. To address this, we performed immunoblotting for H3K27Ac and total H3 levels at different time points after radiation in gliomaspheres. Intriguingly, we saw that the total histone 3 (TH3) levels were reduced at 6 hrs, but significantly increased around 24-48 hrs post-radiation. We also observed an increase in H3K27Ac levels at the same points (Fig 5E, F). To further verify this, we immunostained for total H3 and H3K27Ac in radiated and control tumor sections from orthotopic xenografts. Qualitatively and quantitatively, we found significant reduction in total-H3 positive tumor cells, and on the other hand, a significant increase in AcH3 positive cells post-radiation (Fig 5G-H). We also examined radiated and control mGBM tumors and found significant increase in AcH3 positive tumor cells post-treatment **(Fig S5D).** Finally, we assessed whether the vascular gene regions identified in ATAC-sequencing showed H3K27Ac in radiated gliomaspheres. ChIP-qPCR analysis showed a significant increase in enrichment of genomic regions associated with CDH5 and ANGPT1, but no difference in the housekeeping gene RPL30 in radiated tumor cells immunoprecipitated with anti-H3K27Ac antibody (Fig 5I). Together, these findings indicate that radiation-resistant tumor cells display increased accessibility and H3K27Ac levels in regions associated with vascular genes.

### Blocking P300 histone acetyltransferase activity inhibits radiation-induced vascular-like phenotype switching in glioma cells

Histone acetyltransferases (HAT) are the enzymes that catalyze the transfer of acetyl groups to core histones regulating chromatin structure and gene transcription (32). Since radiation alters H3K27ac in vascular genes, we sought to determine whether blocking histone acetylation prior to radiation inhibits vascular-like phenotype switching in gliomaspheres. First, we examined our RNA-sequencing data to determine which histone acetyltransferases (HAT) are expressed in radiated tumor cells. Of the 14 known HAT enzymes, the EP300 (KAT3B) and KAT2B transcripts showed uniform expression in radiated tumor cells at both 2 and 7-days post-treatment. (Fig 6A). EP300 was also enriched in GEC and GPC **(Fig S6A**). Hence, we decided to target the HAT activity of P300 using a selective small molecule inhibitor, C646 (33). We validated the inhibitor by examining its effect on H3K27ac by immunoblotting (Fig 6B). Quantitative RT-PCR analysis of endothelial and pericyte markers showed significant downregulation in combined C646- and radiation-treated gliomaspheres compared to radiation treatment alone (Fig 6C). This inhibitory effect was seen in two other GBM lines as well as in the murine GBM model **(Fig S6B-D).** We also found significant inhibition of radiation-induced vascular marker expression in CD133+ GSC fraction with C646 treatment (Fig 6D **and Fig S6E).** Furthermore, we utilized the endothelial (CDH5-mCherry) and pericyte (DES-mCherry) reporters and determined that pre-treatment with C646 significantly reduced the expression of the reporters in response to radiation in gliomaspheres, as well as in sorted CD133+ GSC fraction (Fig 6E, F **and Fig S6F, G**).

**Figure 6:**
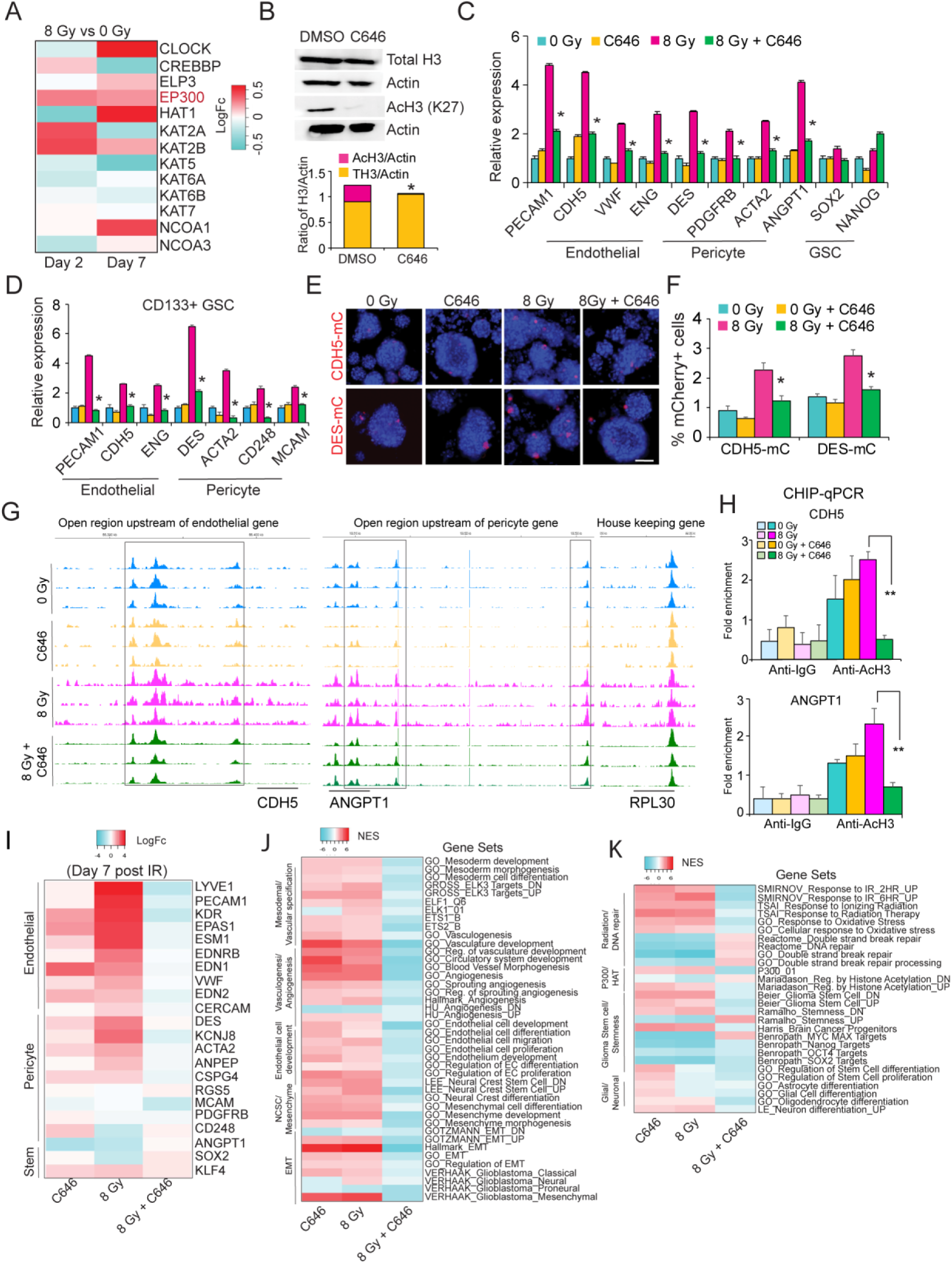
Blocking P300 HAT activity inhibits radiation-induced vascular-like phenotype switching. (A) Heatmap shows logFc expression of HAT transcripts in cultured control and radiated gliomaspheres. EP300 is highlighted in red. (B) Immunoblot images of Total and AcH3 in DMSO and C646 treated gliomaspheres. Quantitation of protein level is shown in the graph. N=2, and * indicates p<0.05, two-tailed t-test. (C, D) Quantitative RT-PCR analysis of endothelial, pericyte and stem genes in control and radiated gliomaspheres and sorted CD133+ GSC fractions treated with C646. Error bars represent mean±SD, N=3, and * indicates p<0.05, two-tailed t-test. (E, F) Images of mCherry expression in control and radiated cells treated with C646. Flow-cytometric quantitation of percentage of mCherry+ cells in each group is shown in the graph. * indicates p<0.05, N=3, two-tailed t-test. (G) Differentially open peak regions (highlighted in black boxes) in vascular genes CDH5, ANGPT1 in control and radiated cells alone or treated with C646. Fold enrichment was normalized to input control. N=2 independent experiments, and ** indicates p<0.005, one-way ANOVA followed by t-test. (H) Graphs show fold enrichment of CDH5 and ANGPT1 genomic regions immunoprecipitated with anti-H2K27Ac and control anti-IgG antibodies. (I) Heatmap of LogFC expression of endothelial, pericyte and stemness genes in control and radiated gliomaspheres treated with and without C646. (J,K) Heatmap shows significant NES (normalized enrichment scores) of gene sets in control and radiated glioma cells treated with and without C646. See also Figure S6.

To determine whether P300 HAT activity is required for radiation-induced chromatin accessibility and H3K27ac of vascular gene regions, we performed ATAC-sequencing and ChIP-qRT-PCR on control and radiated gliomaspheres treated with C646. Surprisingly, we found a significant increase in peak counts, and peak distribution specifically in intergenic regions with combined radiation + C646 treated glioma cells, whereas there were no significant differences amongst control, C646 or radiation treatment alone (N=3 replicates per condition, p<0.05, one-way ANOVA, post-hoc t-test, **Fig S6H, I**). PCA showed clear separation of radiated cells from radiation + C646 treated cells, and from control gliomaspheres (**Fig S6J**). We also found significant differences in number of both up- and down-regulated genes associated with the peaks between radiation + C646 treated cells vs radiation alone (**Fig S6K**), indicating that pre-treatment with P300 HATi can profoundly affect radiation-induced changes in chromatin.

We next examined whether C646 treatment specifically reversed the effects of radiation-induced increase in chromatin accessibility in vascular genes. Indeed, the genomic regions in CDH5 and ANGPT1 that showed increased accessibility (peak size) in radiated cells were reduced with C646 pre-treatment. As expected, GO analysis of differentially open regions showed significant enrichment of terms associated with vascular development, and mesenchymal transition in radiated cells. However, the combined C646 + radiation treatment did not show enrichment of vascular development related terms, but instead showed enrichment of terms related to mitosis, neuron development and chromatin organization and adhesion assembly (**Fig SL**). To demonstrate that P300-mediated H3K27ac is required for radiation-induced changes in chromatin in vascular gene regions, we performed ChIP-qRT-PCR with anti-H3K27ac antibody on control, radiated and C646+ radiation treated gliomaspheres. Pre-treatment with C646 significantly reduced the enrichment of genomic sites in CDH5 and ANGPT1, but not the housekeeping gene RPL30 in radiated cells (Fig 6H), thus confirming that P300 HAT activity is essential for radiation-induced epigenetic rewiring in glioma cells (Fig 6K).

To determine if alterations in chromatin states in vascular genes is reflected in their transcriptional activation, we performed RNA-sequencing in glioma cells treated with 7-day radiation alone or pre-treated with C646 and control cells. In line with the ATAC-sequencing results, treatment with C646 significantly changed gene expression in control and radiated cells, and PCA analysis showed clear separation of radiated and C646+radiation treated cells **(Fig S6M-N).** More importantly, pre-treatment with C646 reduced the expression of several vascular genes induced by radiation (Fig 6I). This is also reflected in GSEA where gene sets related to vascular development, angiogenesis, and mesenchymal transition are diminished, and those associated with DNA repair were enriched (Fig 6J, K). Taken together, these findings clearly established that P300 HAT activity is essential for radiation-induced epigenetic rewiring and vascular-like phenotype switching in glioma cells.

### EP300-deficient glioma cells show reduced vascular-like phenotype and tumor growth post-radiation in vivo

To verify that P300 is essential for radiation-induced vascular phenotype switching *in vivo*, we utilized lentiviral-shRNA to knockdown EP300 in gliomaspheres. The knockdown efficiency was verified by quantitative RT-PCR analysis of EP300 transcript and immunostaining for P300 protein (**Fig S7A, B**). EP300-deficient cells (shEP300) also showed reduced AcH3 compared to control (ShCTL/Scrambled) cells (Fig 7A, B). As expected, and consistent with the C646 inhibitor studies, EP300-deficient cells also showed reduced expression of endothelial and pericyte genes and reporter activation in gliomasphere cultures post-radiation (Fig 7C-E**, Fig S7C)**. Finally, we generated orthotopic tumor xenografts with EP300-deficient and CTL cells and subjected them to radiation. Implantation of EP300-deficient cells showed significant reduction in tumor growth post-treatment and increased animal survival (Fig 7F, G). Immunostaining for total and AcH3 showed fewer AcH3-positive tumor cells with EP300 knockdown, and quantitation revealed that this reduction was significant in radiated, EP300-deficient tumors **(Fig S7D, E).** Immunostaining and quantitation of endothelial (VE-CADHERIN/CD31) and pericyte (DESMIN/aSMA) marker expression in tumor cells (GFP) showed a significant reduction in radiated, EP300-deficient tumors compared to radiated and untreated EP300-deficient or control tumors (Fig 7H, I **and Fig S7F, G).** In summary, these findings indicate that P300 HAT mediates radiation-induced vascular-like phenotype acquisition of glioma cells.

**Figure 7:**
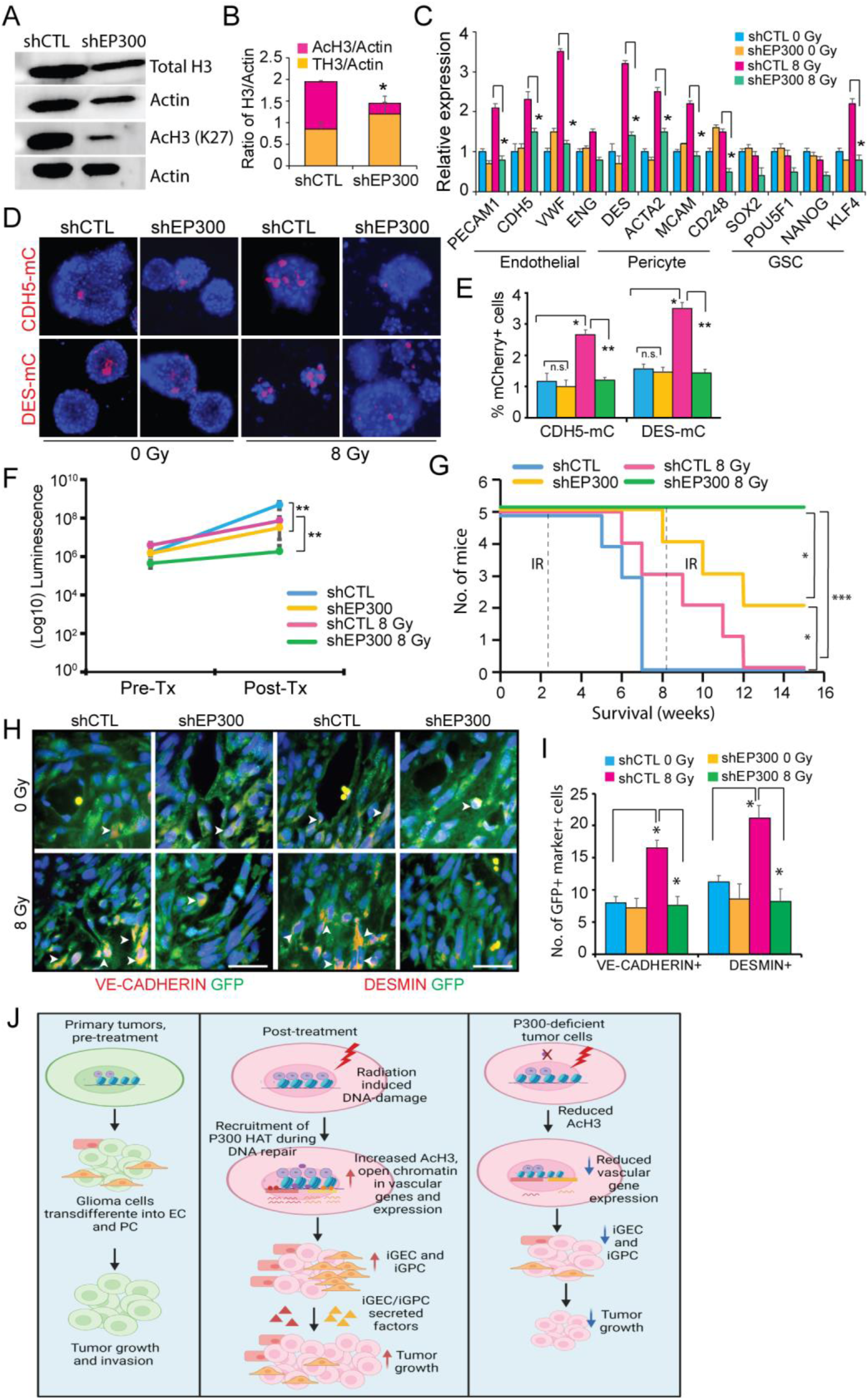
EP300-deficient glioma cells show reduced vascular markers and tumor growth post-treatment. (A, B) Immunoblot images of total and AcH3 in EP300-deficient (shEP300) and control (shCTL) cells. Quantitation of protein levels is shown in the graph. N=2, and * indicates p<0.05, two-tailed t-test. (C) Quantitative RT-PCR analysis of endothelial, pericyte and stem genes in EP300 deficient and control cells post-radiation. N=3, and * indicates p<0.05, two-tailed t-test. (D, E) Images show mCherry expression in EP300-deficient and control cells post-radiation. Flow-cytometric quantitation of percentage of mCherry+ cells in each group is shown in the graph. N=3, * indicates p<0.05, two-tailed t-test. (F) Graph shows quantitation of tumor growth pre- (Pre-Tx) and post-radiation (Post-Tx) treatment. N=5 mice per group, and ** indicates p<0.005, one-way ANOVA, post-hoc t-test. (G) Kaplan-meier survival curve of mice pre- (Pre-Tx) and post-radiation treatment (Post-Tx). * and *** indicates p<0.05, and p<0.0005, Log-rank t-test. (H,I) Immunostaining images VE-CADHERIN and DESMIN in GFP+ tumor cells. Arrows point to marker+ GFP+ cells in each group. Scale bars, 50μm. Quantitation of GFP+marker+ cells in tumor mass is shown in the graph. N=5 mice per group. Error bars represent mean±SD, and * indicates p <0.05, two-tailed t-test. See also Figure S7. (J) Model illustrating radiation-induced vascular-like phenotype switching in glioma stem- and tumor cells via a p300-dependent mechanism.

## Discussion

Despite remarkable progress in understanding the molecular underpinnings of glioma, and the proven benefits of radiation and TMZ, GBM virtually always recur, leading to patient death. A major challenge in developing targeted therapies for GBM is the molecular heterogeneity and treatment-induced phenotypic plasticity exhibited by GSC leading to aggressive recurrence and therapeutic resistance. Elucidating the cellular and molecular mechanisms driving this plasticity can have profound implications for developing new treatments for GBM.

In this study, we show at the single-cell level that radiation significantly alters the functional states of glioma cells. Primarily, radiation induces the phenotypic transition of GSC and tumor cells into vascular endothelial-like and pericyte-like cells, which in turn provide trophic support to the radiated tumor cells and promote recurrence. Radiation induces the vascular-like phenotype switch by altering the accessibility of the chromatin in specific vascular gene regions leading to their increasing expression. This phenotypic switch can be blocked by inhibiting the HAT activity of P300, which highlights a key role for HAT in regulating treatment-induced plasticity, and a novel avenue for targeting and preventing tumor relapse in GBM.

Prior studies have indicated that glioma-derived endothelial cells, while relatively rare, can incorporate into the vasculature, although their role in carrying blood has not been proven (12,26). Glioma-derived pericytes are more common and their functional roles in glioma progression have been more clearly defined (15,34). Our findings indicate that while radiation markedly enhances the frequency of GEC and GPC, these cells do not appear to incorporate into the vasculature to any great extent to play an important role in carrying blood supply. Rather, our data suggest that these cells provide the remaining tumor cells with trophic support, allowing them to survive the severe stress of radiation. The concept of a “vascular niche” that provides trophic support is a common finding in stem cell biology. In the mammalian central nervous system, endothelial cells provide a niche that supports the survival and self-renewal capacity of neural stem cells (35). Studies in rodent models indicate that the vascular niche allows tumor cells to survive radiation-induced stress (36,37). Our results indicate that radiation may cause gliomas to, in fact, create their own supportive niche. The factor or factors elaborated by the radiation-induced GEC and GPC are yet to be elucidated, but will undoubtedly be the targets for potential therapeutic intervention for GBM relapse.

Several independent studies have reported that GSC conversion to endothelial-like cells in tumors is mediated by factors such as HIF1A, NOTCH1, ETV2, WNT5A, TIE2 signaling, whereas pericyte-conversion is driven by TGF-β signaling (12,13,15,26,27,38,39). Our transcriptomic analyses did not reveal significant enrichment of many of these factors in either endothelial-like or pericyte-like clusters or tumor cells post-radiation. This indicated that radiation-induces vascular-like phenotype switching in GSC may occur via a different molecular mechanism. Radiation has been shown to alter histone gene expression and induce methylation changes in cell lines and in animal studies (40,41). Because cell state transitions and phenotype switching requires alterations in the epigenome, we posited that epigenetic rewiring during DNA repair following radiation-induced damage may promote phenotype plasticity in glioma cells. Although our ATAC-sequencing results of radiated gliomasphere cultures did not show significant alterations in global chromatin accessibility due to a high degree of variability between the peak counts obtained from independent experiments, we did however, find that the chromatin was markedly altered in vascular gene regions in radiated cells in all the samples. GO analysis of differentially open regions also revealed enrichment of terms associated with vascular development, endothelial differentiation and mesenchymal transition, supporting our hypothesis that chromatin rewiring in specific vascular genes by radiation drives vascular-like phenotype switching in glioma cells.

Lysine acetylation of histones is a key post-translational modification that regulates chromatin accessibility and gene expression (30). We found increased levels of H3K27ac in vascular gene regions in radiated cells, which indicated that blocking histone acetylation may inhibit the phenotype switch induced by radiation. Histone acetyltransferases (HAT) are the enzymes that catalyze the transfer of acetyl groups onto core histones resulting in altered chromatin structure (32). Of all the HAT genes, P300/KAT3B was enriched in radiated tumor cells and vascular-like cells. It is well established that P300 plays a critical role in the DNA damage response by facilitating repair at sites of double strand breaks (DSB) through acetylation of histones and chromatin decompaction (42–45). Selective inhibition of P300 HAT activity using C646 small molecule inhibitor has been shown to inhibit cell growth and sensitize cells to DNA damaging agents in other cancers including melanoma (46) NSLC (47), colorectal (48), prostrate (49,50) and neuroblastoma (51). Consistent with these studies, we found that blocking P300 HAT activity resulted in significant reduction of chromatin accessibility and H3K27ac in vascular gene regions in response to radiation. However, these effects were highly site-specific to vascular genes induced by radiation as global chromatin accessibility counterintuitively increased with the combined C646 and radiation treatment. RNA-sequencing results also mirrored the ATAC-sequencing data as C646 treatment reduced vascular gene expression and gene sets associated with angiogenesis induced by radiation, but conversely increased metabolic and cellular biosynthesis-related processes that decreased with radiation. These results suggest that P300 HAT activity is selectively required for radiation-induced epigenetic changes at sites of vascular genes. But its disruption might also dysregulate the global DNA repair response and chromatin integrity. Further studies will be needed to determine the precise mechanism by which P300 mediates chromatin decompaction and DNA repair in radiated glioma cells.

The net functional outcome of P300 disruption in radiated tumor cells is the inhibition of the acquisition of vascular-like phenotypes, reduction of tumor growth and enhanced animal survival, indicating that P300 is a potential target for enhancing the effects of radiation. P300 may also play roles in promoting GBM growth outside of the context of radiation. Prior studies have suggested that P300 can be either tumor suppressive or oncogenic (52). In our GBM models, we find that P300 plays a pro-tumorigenic role, even in the absence of radiation, as P300-deficient tumors diminish tumor growth *in vivo*. This effect is unlikely to be mediated by an effect on the production of vascular-like cells by non-irradiated GSC, as we did not observe any significant impact of P300 inhibition or knockdown on vascular gene expression. Further studies will be needed to assess the general role of P300 in gliomagenesis.

The findings presented here provide new avenues to address radiation-induced cellular plasticity and resulting resistance in GBM. Radiation-induced GEC and GPC play important trophic roles in promoting the survival of irradiated GBM cells, and thus preventing the acquisition of these cellular states could enhance the therapeutic response to radiation and potentially prevent or delay tumor recurrence. Whether similar processes occur in other radiation-treated cancers remains to be seen. Furthermore, our findings highlight the importance of investigating HAT as key epigenetic drivers mediating pro-tumorigenic programs and their broad applicability for treating cancer.

## Methods

### 1. Patient-derived gliomasphere cultures

All patient-derived gliomasphere lines were cultured and maintained as previously described (53). Experiments were performed only with lines that were cultured for < 20 passages since their initial establishment, and tested negative for mycoplasma contamination.

### 2. Irradiation of gliomaspheres and animals

Cultured cells were irradiated at room temperature using an experimental X-ray irradiator (Gulmay Medical Inc. Atlanta, GA) at a dose rate of 5.519 Gy/min for the time required to apply a prescribed dose. The X-ray beam was operated at 300 kV and hardened using a 4mm Be, a 3mm Al, and a 1.5mm Cu filter and calibrated using NIST-traceable dosimetry. Tumor-bearing mice were irradiated at a single dose of 8 Gy using an image-guided small animal irradiator (X-RAD SmART, Precision X-Ray, North Branford, CT) with an integrated cone beam CT (60 kVp, 1 mA) and a bioluminescence-imaging unit as described previously (Bhat et al. 2020). Individual treatment plans were calculated for each animal using the SmART-Plan treatment planning software (Precision X-Ray). Radiation treatment was applied using a 5×5mm collimator from a lateral field.

### 3. Animal strains, intracranial transplantation, treatments and imaging

All animal studies were performed according to approved protocols by the institutional animal care and use committee at UCLA. Studies did not discriminate sex, and both male and females were used. **Strains**: 8 to 12-week old NOD-SCID gamma null (NSG) mice were used to generate tumors from a patient-derived GBM line HK_408, and a murine IDH1-wt GBM model (wild-type IDH1 (NRAS G12V-shp53-shATRX) described previously (22). 5X10^4^ tumor cells containing a firefly-luciferase-GFP lentiviral construct were injected intracranially into the neostriatum in mice. For assessing tumor initiation potential of CD133+ and CD133-fractions, 1X10^3^ cells injected intracranially. Co-transplantation experiments were performed at a ratio of 1:1 (tumor: vascular-like cells) 5X10^4^ cells per condition. **Treatments:** For GCV treatment, animals were injected by i.p at a dose of 80mg/kg of body weight of mice every day for a week, and for DT treatment, animals were given i.p injections at a dose of 5μg/kg of body weight of mice every 2 days over a week. **Imaging**: Tumor growth was monitored 2 weeks after transplantation by measuring luciferase activity using IVIS Lumina II bioluminescence imaging. ROIs were selected to encompass the tumor area and radiance was used as a measure of tumor burden.

### 4. Single-cell RNA sequencing and analysis

Detailed protocol can be found in the supplemental information

### 5. Bulk RNA-sequencing and analysis

Detailed protocol can be found in the supplemental information

### 6. ATAC-sequencing and analysis

Detailed protocol can be found in the supplemental information

### 7. Chromatin Immunoprecipitation-quantitative RT-PCR

ChIP was performed on 5X10^6^ cells from 2-day control and radiated gliomaspheres using SimpleChIP® Plus Enzymatic Chromatin IP Kit (Magnetic Beads) (Cell Signaling Technology, #9005S) according to manufacturer’s protocol. AcH3-K27 and Normal IgG antibodies (Cell signaling, #8173S and #2729) were used for immunoprecipitation, and primers for qRT-PCR to examine amplification of vascular genes regions are listed in supplementary information.

### 8. Statistical analysis

All data are expressed as the mean ± SD. Quantification of qRT-PCR, cell-based assays, and flow cytometry are representative of at least three independent experiments unless otherwise stated. *P* values were calculated in Graph Pad Prism 8.0 using Welch’s t-test or unpaired two-tailed Student t-test and ANOVA for multiple comparison, followed by post-hoc t-test which can be found in the individual figures and figure legends. *P* values less than 0.05 were considered to be significant. Log-rank analysis was used to determine the statistical significance of Kaplan-Meier survival curves, and no samples, mice or data points were excluded from the analysis reported in this study. R-package was used for statistical analysis of all sequencing experiments.

See supplementary information for details and other methods.

## Financial support

Broad Stem cell postdoctoral fellowship (SDM), UCLA. The Dr. Miriam and Sheldon G. Adelson Medical Research Foundation (RK, SAG, DHG, HIK), the UCLA SPORE in Brain Cancer P50 CA211015-01A1 (FP, HIK), NIH R01HL149687 (AD), NIH R01CA200234 (FP) and the UCLA Intellectual and Developmental Disability Research Center P50 HD103557 (HIK).

## Acknowledgements

The authors thank the UCLA pathology, flow cytometry, TCGB and UNGC sequencing cores for their technical assistance with support from the Jonsson Comprehensive Cancer Center P30CA016042 and Dr. Paul Mischel for helpful discussions on the manuscript.

## Author contributions

S.M and H.K conceptualized, designed the experiments and wrote the manuscript. S.M, P.N, R.P, M.J, N.V, M.C, A.A, R.G, A.P. and Q.W conducted the experiments. R.K and Y.Q. performed the bioinformatics analysis. T.B, D.G, M.C, and P.L contributed resources, reagents and contributed to experimental discussions. F.P, A.D, J.H, and S.G contributed valuable inputs in designing experiments and data analysis.

## Conflict of Interest

The authors declare no competing interests.

## Notes

### Competing Interest Statement

The authors have declared no competing interest.

### Summary of Updates

The manuscript has been updated with the following changes: a. Revised title, and abstract. b. Results and discussion sections have been revised to include some new data on radiation-induced changes in chromatin accessibility and the role of P300 HAT in regulating H3K27ac and chromatin state in vascular gene regions. Figures 5 and 6 and their corresponding supplemental figures are also revised. c. Supplemental files are also updated.

