## Supplementary Information for "Radiation-reprogrammed glioma stem cells generate vascular-like cells to build a trophic niche driving tumor recurrence"

**Supplemental Information**

**Materials and Methods:**

1. **Single-cell RNA sequencing and analysis:** Preparation of single cell suspensions: **Gliomaspheres:** Single cell suspensions of cultured control (0 Gy) and 2- and 7-day radiated (8 Gy) gliomaspheres were prepared on the same day using 10x Genomics Chromium Single Cell 3' Reagent Kits v3 according to the manufacturer's protocol. **Tumor xenografts**: Control and radiated tumors (N=3) were enzymatically dissociated with Collagenase II, Trypsin and DNAse to generate cell suspensions. Contaminating mouse cells were removed using MACS mouse cell depletion kit (Miltenyi Biotec). Viable GFP+ tumor cells were FACS sorted and collected for preparing single cell suspensions using 10x Genomics Chromium Single Cell 3' Reagent Kits v3. Quality control and sequencing: Quantity and quality of cDNA were assessed by Agilent 2100 expert High Sensitivity DNA Assay. cDNA samples were sequenced on 1 lane of NovaSeq 6000 S2 flowcell. Reads were mapped to HumanGRCh38 genome using Cell Ranger v.3.0.2. **Gliomaspheres:** More than 600 million reads were obtained for each sample. Average number of genes detected was 5875 (SE±357). Confident read mapping rates were 81.2-87.4% with over 86.8% of reads in cells**.** Filtering genes and cells: Seurat package v3.1.1 was used to do analysis. For each condition, genes expressing in less than 5 cells were removed. Cells with number of features less than 500 were excluded. PCA was used for dimension reduction with top 5000 most variable genes. Raw count data were normalized using regularized negative binomial regression with SCTransformation, batch correction was done with Seurat's integration method using Canonical Correlation Analysis (CCA) with 75 dimensions. Cell clustering is done using Shared Nearest Neighbor (SNN) Graph method. Markers were identified by Wilcox Rank Sum test. Directional p-Values of the entire gene sets were used for enrichment analysis by GSEA. Each cell cluster was annotated by a combination of the following methods 1) canonical marker expression 2) Gene Set Enrichment Analysis (GSEA) of cluster specific markers determined by Find Markers function in Seurat and 3) Co-

expression modules obtained by Louvain community detection clustering method. For GSEA, MSigDB ver7.0 was used as reference. Trajectory analysis was done with monocle 3 (version 0.2.1.3) by converting Seurat object to monocle. **Tumor xenografts:** Over 200 million reads were obtained for each sample. Average number of genes detected was 6362 (GBM_TX 0 Gy) and 4824 (GBM_TX 8 Gy), and confident read mapping rates were over 76.5-78% with over 88% of reads in cells. *In vivo* single cell dataset was similarly analyzed to *in vitro* dataset except following: the two *in vivo* expression matrices from CellRanger output were merged, then low expression genes present in less than 5 cells and cells with less than 500 or more than 8000 features, as well as mitochondrial contents greater than 5 percent were filtered. Mitochondrial and ribosomal genes were excluded for downstream analysis. Filtered expression data were individually normalized using SCTransform in Seurat functions. PCA was used for dimension reduction with top 3000 most variable genes. Batch effect was corrected using harmony.

2. **Bulk RNA-sequencing and analysis:** RNA extraction from gliomaspheres (3 replicates per condition) was performed using Qiagen RNeasy microkit. RNA quality was assessed using Bioanalyzer and only samples with a RIN score >8.0 were sequenced. RNA samples were pooled and barcoded, and libraries were prepared using TruSeq Stranded RNA (100ng) + Ribozero Gold. Paired-end 150bp reads were aligned to the latest human_hg38 reference genome using the STAR spliced read aligner. Total counts of read-fragments aligned to known gene regions within the human hg38 refSeq reference annotation was used as the basis for quantification of gene expression. Differentially expressed genes were identified using DESeq and edgeR, which are then considered and ranked based on False Discovery Rate (FDR Bejamini Hochberg adjusted p-values of ≤ 0.01). Gene Set Enrichment Analysis (GSEA) was carried out to determine the gene signatures differentially regulated in control and radiated cellsand represented as heatmaps.

3. **Quantitative RT-PCR:** RNA was isolated with RNeasy Micro or Mini Kit (QIAGEN), and then used for first-strand cDNA synthesis using random primers and Superscript Reverse Transcriptase (Invitrogen). qRT–PCR was performed using Power SYBR Green PCR Master Mix (Applied Biosystems). The relative expression of genes was normalized using 18srRNA as the housekeeping gene. All experiments were repeated at least 3 times, and data is represented as mean ± SD. Primers are listed in the Table S4.

**4. Lentivirus-transduction in tumor cells:** Lentiviral vectors (CDH5-mCherry, DES-mCherry, DES-GFP, CDH5-HSV-TK-mCherry, DES-HSV-TK-mCherry, CDH5-DTR-mCherry and DES-DTR-mCherry) were designed and purchased from Vector builder. EP300 and Scrambled-siRNAs were purchased from Abmgood. Gliomaspheres were transduced with indicated viruses for 48 hours and selected with puromycin (Sigma). Reporter gene expression and knockdown was validated by immunofluorescence, quantitative RT-PCR or immunoblotting in target cells.

**5. Flow cytometry and FACS sorting:** Cells were harvested and suspended in ice-cold PBS with 1% BSA and 2mM EDTA. After incubation with FcR Blocking Reagent (Miltenyi Biotec), cells were stained by fluorescently conjugated antibodies and incubated for 10 min in the dark in the refrigerator (2−8°C). Antibodies include CD31-PE, CD144-APC, CD146-PE CD248-APC, IgG-APC and IgG-PE from Miltenyi Biotec. The stained cells or GFP and mCherry-labeled cells were analyzed in a BD Fortessa analyzer. FACS sorting was performed using the BD FACSAria cell sorter. Data were analyzed using FlowJo software.

6. **Immunofluorescence (IF) staining and immunoblotting:** For IF staining, 5μm FFPE brain sections were incubated with primary antibodies overnight at 4°C after deparaffinization,

rehydration, antigen retrieval and blocking in PBS with 2% BSA. Sections were then incubated with species-appropriate goat/donkey secondary antibodies coupled to AlexaFluor dyes (488, 568 or 647, Invitrogen) and Hoechst dye for nuclear staining for 2 hrs at RT. VECTASHIELD (Vector Laboratories) was used to mount coverslips. Slides were imaged using Leica LAS X or EVOS microscope, and quantification was performed manually using ImageJ. For quantitation of stained sections, marker+ GFP+ tumor cells in the tumor mass or blood vessels spanning 200μm area/section was counted in 10 random sections per condition, and data is represented as mean ± SD in the graphs.

For **immunoblotting**, cells were harvested, washed and lysed in RIPA buffer with protease inhibitor cocktail and centrifuged at 13,000 rpm for 10 min. Protein concentration in each sample was determined by Bradford assay using BSA as a standard. 15ug of protein was used for each sample. Protein lysates were subjected to SDS-PAGE on 4%–12% gradient polyacrylamide gel (Thermo-fischer Scientific), transferred onto nitrocellulose membranes and incubated with primary antibodies, washed, and probed with HRP-conjugated secondary antibodies. Quantitation of protein levels was performed in ImageJ by normalizing to loading control, b-actin. Data is represented as mean ± SD, and p-values are derived from two independent experiments. Antibodies are listed in Table S3.

**7. Matrigel tube formation and matrigel plug in vivo angiogenesis assay:** Tubular network formation was assessed by growth factor reduced (GFR) Matrigel assay kit (BD Biosciences) in three-dimensional (3D) culture according to the manufacturer's instructions. Briefly, tumor cells were harvested 7days post-radiation and cultured on GFR matrigel for 16-24 hours. Images were obtained at 20x magnification on EVOS microscope andnumber of branch points was quantitated manually from 5 random fields per well and 3 replicates per group. **Matrigel plug in vivo angiogenesis assay:** $5X10^5$ labelled sorted GFP+ tumor and mCherry+ GFP+ transdifferentiated cells from control and radiated groups were embedded in

GFR matrigel. Matrigel containing cells were injected subcutaneously into mice and allowed to solidify to form a plug. 7 days after implantation, plugs were harvested and fixed in 4% PFA. Sections were stained with Hematoxylin & Eosin stain and Masson's Trichrome to label the basement membrane and blood vessels. Images were obtained at 20x magnification using EVOS microscope. Quantitation of vessel density was performed using Image J plugin for Vessel analysis. Percent vessel density in each condition represents mean ± SD derived from 3 technical replicates and from 3 independent experiments.

8. **DiI-Ac-LDL uptake assay:** Control and radiated gliomasphere cultures were incubated with DiI-Ac-LDL for 24 hours. Cells were fixed, and stained with VE-CADHERIN and imaged using EVOS microscope. Quantitation of number of DiI-Ac-LDL+ VE-CADHERIN+ cells was performed by manually counting the double-positive cells from 10 random fields per condition. Data is represented as mean ± SD in the graphs, and p-values are derived from 3 independent experiments.

**9. Limiting dilution assay:** For assessment of self-renewal of sorted CD133+ and CD133- fractions, cells were plated at a density of 1, 5, 10, 25 and 50 cells per well (10 wells for each density). Cells were maintained for 10 days before sphere formation was evaluated. Spheres larger than 10 cells in diameter were considered for analysis. Numbers represent stem cell frequency as calculated using the Walter and Eliza Hall Institute Bioinformatics Division ELDA analyzer http://bioinf.wehi.edu.au/software/elda/

**10. Cell proliferation assay:** GBM cells were plated at a density of 5000 cells per well in 96-well plates. Proliferation was assessed 3 days after treatment with inhibitors or conditioned media using CellTiter-Glo® Luminescent Cell Viability Assay. Luminescence signal was measured in a luminometer, and readings were taken on Day 0 of plating and at Day 3 after treatment to normalize for plating density and obtain fold change in growth of cells.

**11. ATAC-sequencing and analysis:** Chromatin accessibility of control and radiated gliomaspheres (3 replicates per group) was performed as described previously (1). ATAC-Libraries from each condition was sequenced on NextSeq 500 High Output Kit v2 (150 cycle, FC-404-2002, Illumina). Alignment of reads was carried out using the Burrows-Wheeler Aligner mem using hg19 assembly. Peak calling was performed using MACS2 (with parameter setting –nomodel –shift 75), and differential peak analysis using featureCount and DESeq2 (default setting). Motif analysis and peak annotation was done using HOMER and GO analysis using HOMER and EnrichR. UCSC Genome Browser was used to determine whether open regions displayed H3K27Ac and conserved TF binding sites. Integrated Genome viewer (IGV) was used to represent the peaks/open regions.

**12. Chromatin Immunoprecipitation (ChIP)-quantitative RT-PCR:** Chromatin immunoprecipitations were performed using SimpleChIP® Enzymatic Chromatin IP Kit (Magnetic beads, Cell Signaling Technology, #9005S #) according to manufacturer's instructions. Briefly, formaldehyde cross-linked chromatin from each sample is enzymatically digested using Micrococcal Nuclease, followed by sonication to obtain chromatin fragments. Fragmented chromatin was incubated with anti-rabbit IgG or anti-rabbit H3K27Ac antibodies for immunoprecipitation with Protein-G magnetic beads. After reverse-crosslinking, DNA is purified and analyzed by quantitative RT-PCR. Primers for human RPL30 was provided with the kit. Primers for CDH5 and ANGPT1 were designed specifically for open regions identified from ATAC-sequencing in radiated cells. CDH5 Forward primer: CATAAAAGTCCTTCCCATGTTGC, CDH5 Reverse primer: TGGCAATGAAGAGTAGTCCCAA. ANGPT1 Forward primer: CGACAGTTGCCATCGTGTTC and ANGPT1 Reverse primer: TTTCCTCGCTGCCATTCTGA were custom synthesized from Integrated DNA technologies. Quantitation of DNA was performed using RT--PCR and fold enrichment was normalized to input control.

#### Supplementary Tables

**Table S1:** Key Resources

| Reagents | Source | Identifier/Catalogue number |
|---|---|---|
| **Antibodies** | | |
| Mouse monoclonal anti-CD31, JC70A, human | Agilent DAKO | M082329-2 |
| Mouse monoclonal anti-CD144/VE-CADHERIN, Clone BV9, human | Biolegend | 348502 |
| Mouse monoclonal anti-DESMIN, Clone D33 human | Agilent DAKO | M076001-2 |
| Mouse monoclonal anti-aSMA, Clone 1A4 human | Millipore Sigma | A2547 |
| Polyclonal Goat anti-VE-CADHERIN, human | R&D systems | AF938 |
| Mouse monoclonal anti-VE-CADHERIN, Clone BV9, human | Biolegend | 348502 |
| Rabbit polyclonal anti-GFP | Novus Biologicals | NB600-308 |
| Mouse monoclonal anti-mCherry | Novus Biologicals | NBP1-96752 |
| Chicken polyclonal anti-GFP | Novus Biologicals | NB100-1614 |
| Chicken polyclonal anti-mCherry | Millipore Sigma | AB356481 |
| CD31-PE (WM59), human | Biolegend | 303105 |
| CD144-APC (REA199), human | Miltenyi Biotec | 130-100-708 |
| CD146-APC, Clone P1H12, human | Biolegend | 361015 |
| CD248-647, Clone B1/35 human | BD Pharmingen | 564994 |
| CD133/2-PE, clone 293C3 human | Miltenyi Biotec | 130-113-186 |
| Rabbit monoclonal anti-B-actin | Cell Signaling Technology | 4970S |
| Rabbit polyclonal anti-Histone 3 (D1H2) | Cell Signaling Technology | 4499 |

| | | |
|---|---|---|
| Rabbit polyclonal anti-AcH3 (K27) | Cell Signaling Technology | 4353 |
| Rabbit polyclonal anti-GFAP | Agilent DAKO | GA52461-2 |
| Mouse monoclonal anti-NESTIN, Clone 10C2 human | Millipore Sigma | MAB5326 |
| Rabbit monoclonal anti-VIMENTIN (D21H3) | Cell Signaling Technology | 5741S |
| Rabbit monoclonal anti-N-CADHERIN (D4R1H) | Cell Signaling Technology | 13116S |
| Rabbit monoclonal anti-SOX2 (D9B8N) | Cell Signaling Technology | 23064S |
| Rabbit monoclonal anti-P300 (D8Z4E) | Cell Signaling Technology | 86377S |
| Rabbit monoclonal, anti-DESMIN (D93F5) | Cell Signaling Technology | 5332S |
| Rabbit monoclonal, anti-VE-CADHERIN | Cell Signaling Technology | 2500S |
| Goat polyclonal anti-VE-CADHERIN, mouse | R&D Systems | AF1002 |
| Tomato Lectin-DyLight 649 | Vector Laboratories | DL-1178-1 |
| **Chemicals, reagents and commercial assays** | | |
| Ganciclovir (GCV) | Sellechckem | S1878 |
| Diphtheria toxin (DT) | EMD Millipore Sigma | 322326 |
| C646 | Sellechckem | S7152 |
| Matrigel GFR membrane matrix | Fisher Scientific, Corning | CB40230C |
| CellTiterGlo 2.0 assay | Fisher Scientific | PRG9242 |
| DiI-Ac-LDL Kit | Cell applications | 022k |
| Mouse cell depletion kit | Miltenyi Biotec | 130-104-694 |
| **Experimental Models: Cell Lines** | | |
| Patient-derived gliomasphere lines | This lab | N/A |
| Human Umbilical vein endothelial cells (HUVEC) | Sciencell | 8000 |
| Murine GBM line | | Laboratory of Dr. Maria Castro |
| **Mouse strains** | | |
| NOD.Cg-*Prkdc*$^{scid}$ *Il2rg*$^{tm1Wjl}$/SzJ (NSG) | Jackson Laboratory | 00557 |
| C57BL6 | Jackson Laboratory | 000664 |

| | | |
|---|---|---|
| NRAS G12V-shp53-shATRX-IDH1 wildtype (mGBM model) | Castro Lab | |
| **Lentiviral vectors** | | |
| pLV(EXP)-hDES-mCherry:T2A:puro | Vector Builder | Promoter region: 1786 bp |
| pLV(EXP)-hCD144/CDH5-mCherry:T2A:puro | Vector Builder | Promoter region: -1400 to +100 bp |
| pLV(EXP)-puro-hDES-GFP | Vector Builder | |
| pLV(EXP) - mCherry:T2A:puro-hCD144-DeltaTK | Vector Builder | |
| pLV(EXP) - mCherry:T2A:puro-hDES-DeltaTK | Vector Builder | |
| pLV(EXP) - mCherry:T2A:puro-hCD144-DTR | Vector Builder | |
| pLV(EXP) - mCherry:T2A:puro-hDES-DTR | Vector Builder | |
| **Softwares** | | |
| EVOS FL Auto2 Cell Imaging | Thermofisher Scientific | |
| Leica LAS-X | Leica microsystems | |
| Graphpad Prism 8 | Graphpad | https://www.graphpad.com/scientific-software/prism/ |
| Image J | National Institute of Health | https://imagej.nih.gov/ij/ |
| R-Package V.3.2.5 | The R project for Statistical computing | https://www.r-project.org/ |
| Cell Ranger v 3.0.2 | 10x Genomics | https://support.10xgenomics.com/single-cell-gene-expression/software/ |
| Seurat v. 3.1.1 | Satija Lab | https://satijalab.org/seurat/ |
| FlowJo | FlowJo | https://www.flowjo.com/ |
| GSEA | Broad Institute | http://software.broadinstitute.org/gsea/index.jsp |
| HOMER | Benner/Glass Labs, UCSD | http://homer.ucsd.edu/homer/ |
| Biorender | Biorender | https://biorender.com/ |
| EnrichR | Ma’ayan Lab | https://maayanlab.cloud/Enrichr/ |

**Table S2: Primers for qRT-PCR**

| Genes | Forward primer | Reverse primer |
|---|---|---|
| **Human** | | |

| | | |
|---|---|---|
| 18srRNA | GGCCCTGTAATTGGAATGAGTC | CCAAGATCCAACTACGAGCTT |
| PECAM1/CD31 | CCAAGGTGGGATCGTGAGG | TCGGAAGGATAAAACGCGGTC |
| CDH5 | AAGCGTGAGTCGCAAGAATG | TCTCCAGGTTTTCGCCAGTG |
| FLT1/VEGFR1 | GAAAACGCATAATCTGGGACAGT | GCGTGGTGTGCTTATTTGGA |
| VWF | CCGATGCAGCCTTTTCGGA | TCCCCAAGATACACGGAGAGG |
| ENG | CGCCAACCACAACATGCAG | GCTCCACGAAGGATGCCAC |
| EDN1 | AAGGCAACAGACCGTGAAAAT | CGACCTGGTTTGTCTTAGGTG |
| DES | GAGACCATCGCGGCTAAGAAC | GTGTAGGACTGGATCTGGTGT |
| ACTA2 | CTATGAGGGCTATGCCTTGCC | GCTCAGCAGTAGTAACGAAGGA |
| PDGFRB | AGCACCTTCGTTCTGACCTG | TATTCTCCCGTGTCTAGCCCA |
| MCAM/CD146 | AGCTCCGCGTCTACAAAGC | CTACACAGGTAGCGACCTCC |
| CD248 | TGGTGCCAACGTGTGTCTTTT | AGCGATAGCAGTCAGTGATGC |
| ANGPT1 | AGCGCCGAAGTCCAGAAAAC | TACTCTCACGACAGTTGCCAT |
| SOX2 | GCCGAGTGGAAACTTTTGTCG | GGCAGCGTGTACTTATCCTTCT |
| NES | CTGCTACCCTTGAGACACCTG | GGGCTCTGATCTCTGCATCTAC |
| GFAP | CTGCGGCTCGATCAACTCA | TCCAGCGACTCAATCTTCCTC |
| POU5F1/OCT4 | GGGAGATTGATAACTGGTGTGTT | GTGTATATCCCAGGGTGATCCTC |
| NANOG | TTTGTGGGCCTGAAGAAAACT | AGGGCTGTCCTGAATAAGCAG |
| KLF4 | CGGACATCAACGACGTGAG | GACGCCTTCAGCACGAACT |
| CD15/FUT4 | GATCTGCGCGTGTTGGACTA | GAGGGCGACTCGAAGTTCAT |
| CDH1/E-CADHERIN | AAAGGCCCATTTCCTAAAAACCT | TGCGTTCTCTATCCAGAGGCT |
| CDH2/N-CADHERIN | AGCCAACCTTAACTGAGGAGT | GGCAAGTTGATTGGAGGGATG |
| SNAI1 | TCGGAAGCCTAACTACAGCGA | AGATGAGCATTGGCAGCGAG |
| SNAI2 | TGTGACAAGGAATATGTGAGCC | TGAGCCCTCAGATTTGACCTG |
| TWIST1 | GTCCGCAGTCTTACGAGGAG | GCTTGAGGGTCTGAATCTTGCT |

| | | |
|---|---|---|
| BMI1 | GCTGCCAATGGCTCTAATGAA | TGCTGGGCATCGTAAGTATCTT |
| **Mouse** | | |
| *18srRNA* | GTAACCCGTTGAACCCCATT | CCATCCAATCGGTAGTAGCG |
| *Pecam1* | CTGCCAGTCCGAAAATGGAAC | CTTCATCCACCGGGGCTATC |
| *Cdh5* | CACTGCTTTGGGAGCCTTC | GGGGCAGCGATTCATTTTTCT |
| *Eng* | CCCTCTGCCCATTACCCTG | GTAAACGTCACCTCACCCCTT |
| *Edn1* | GCACCGGAGCTGAGAATGG | GTGGCAGAAGTAGACACACTC |
| *Des* | GTGGATGCAGCCACTCTAGC | TTAGCCGCGATGGTCTCATAC |
| *Mcam* | CCCAAACTGGTGTGCGTCTT | GGAAAATCAGTATCTGCCTCTCC |
| *Cd248* | CAACGGGCTGCTATGGATTG | GCAGAGGTAGCCATCGACAG |
| *Pdgfrb* | CAAGAAGCGGCCATGAATCAG | CGGCCCTAGTGAGTTGTTGT |
| *Sox2* | GCGGAGTGGAAACTTTTGTCC | CGGGAAGCGTGTACTTATCCTT |
| *Pou5f1/Oct4* | GGCTTCAGACTTCGCCTCC | AACCTGAGGTCCACAGTATGC |
| *Nanog* | TCTTCCTGGTCCCCACAGTTT | GCAAGAATAGTTCTCGGGATGAA |
| *Klf4* | GTGCCCCGACTAACCGTTG | GTCGTTGAACTCCTCGGTCT |
| *Nestin* | CCCTGAAGTCGAGGAGCTG | CTGCTGCACCTCTAAGCGA |
| *Gfap* | CCCTGGCTCGTGTGGATTT | GACCGATACCACTCCTCTGTC |

Supplementary Figures:

#### Figure S1

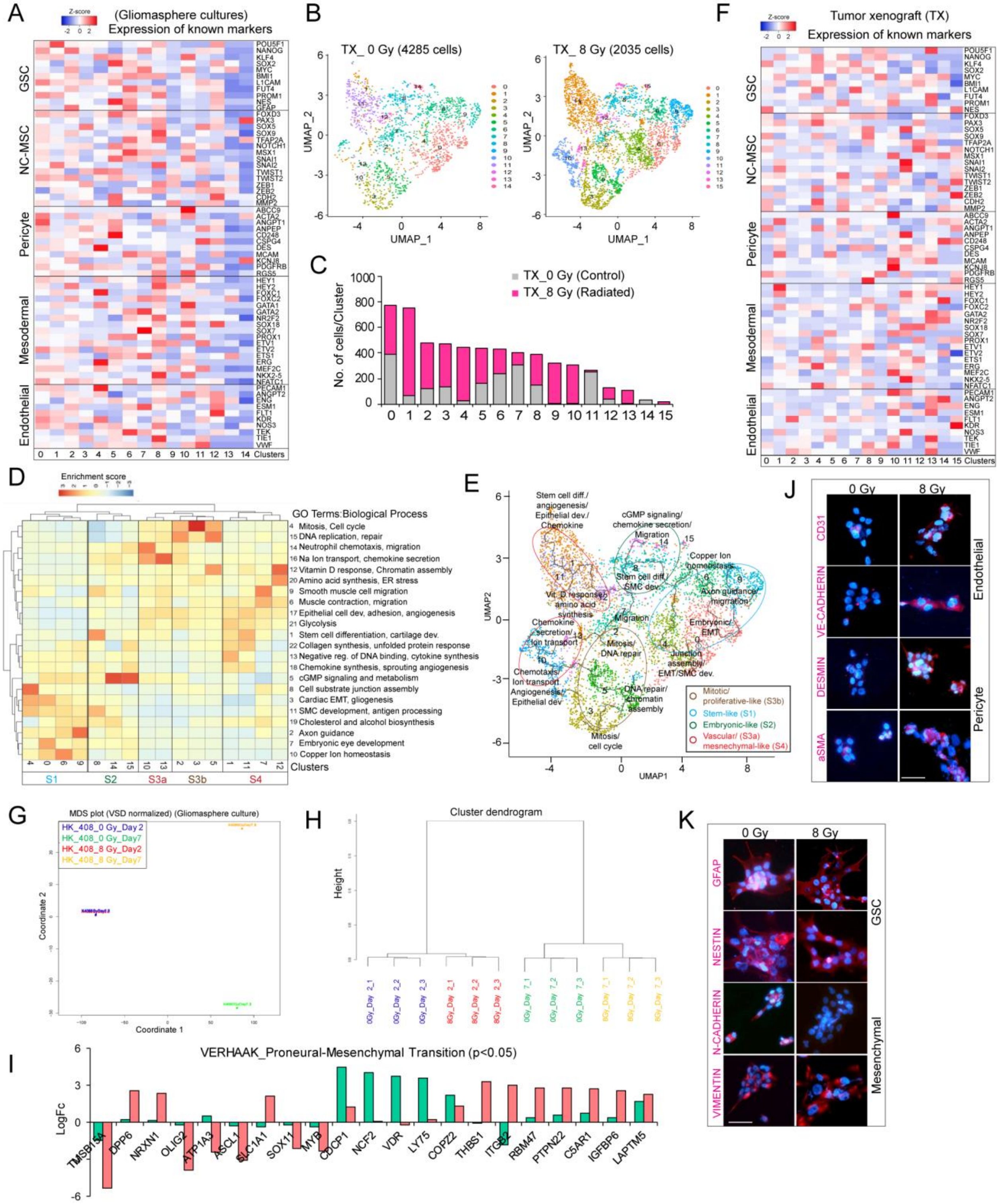

**Figure S1: Single-cell sequencing shows diverse functional states of glioma cells**

(A) Heatmap shows average expression of stemness, neural crest/mesenchymal stem cell (NC-MSC), mesodermal and mature endothelial and pericyte markers in each cluster in cultured control and radiated gliomaspheres.

(B) UMAP plot of glioma cells from control (0 Gy) and radiated (8 Gy) tumor xenografts (TX).

(C) Histogram shows the number of cells per cluster in each group.

(D) Heatmap of differential expression of co-expressed gene modules and Louvain clustering. The subgroups are highlighted in black boxes.

(E) UMAP plot of pseudotime trajectory analysis of all the clusters. The functional states representing the subgroups (circled in color) are shown in the inset.

(F) Heatmap shows average expression of markers in each cluster from tumor xenografts.

(G, H) MDS plot and cluster dendrogram of gene expression differences between control and day 2 and day 7-radiated gliomaspheres.

(I) LogFC values of genes associated with VERHAAK_Glioblastoma_Mesenchymal gene set in control and radiated gliomaspheres.

(J, K) Immunostaining of endothelial, pericyte, GSC and mesenchymal markers in control and radiated gliomaspheres. Scale bars, 100um.

**Figure S2**

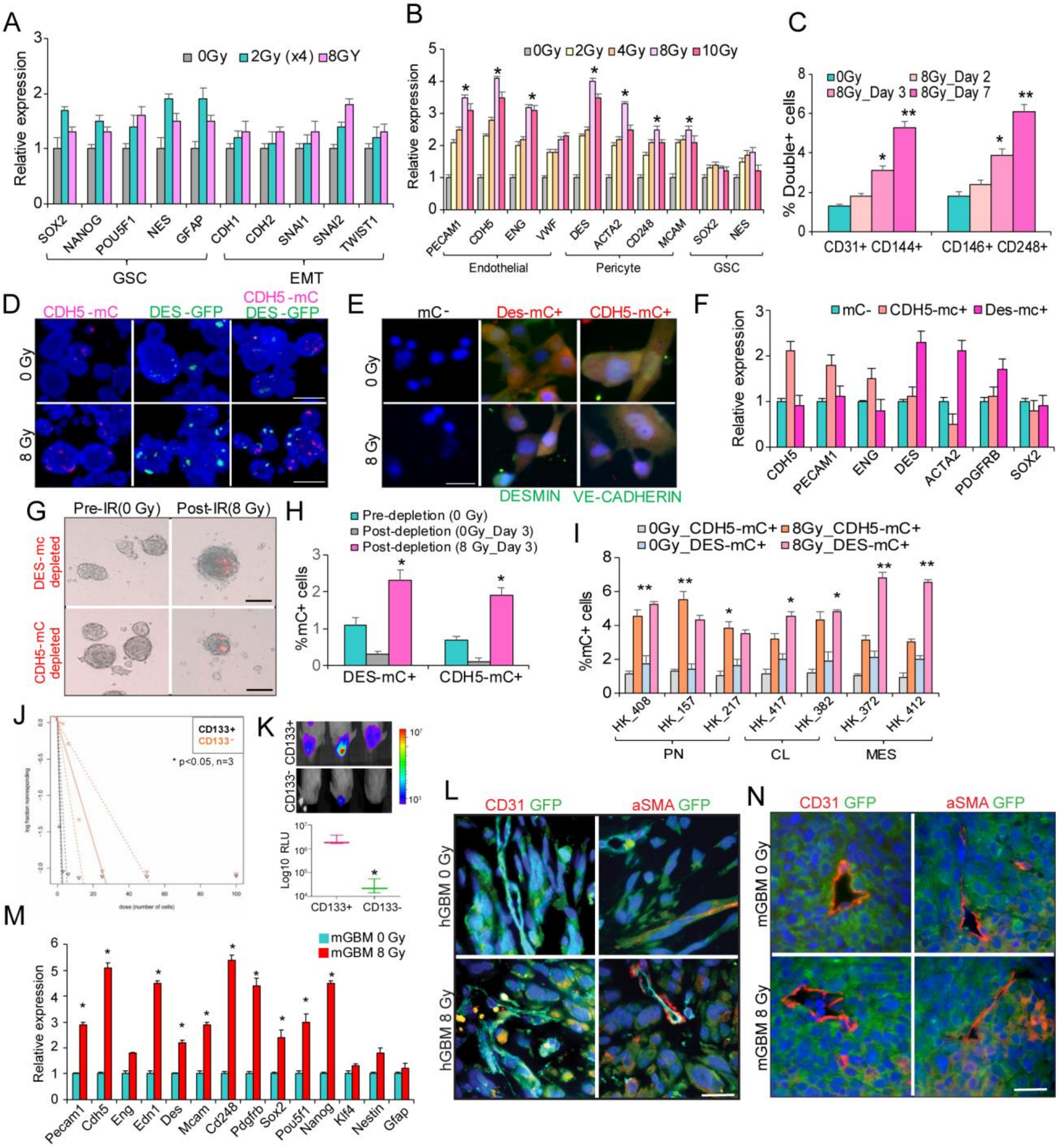
A
0Gy 2Gy (x4) 8GY
Relative expression
SOX2 NANOG POU5F1 NES GFAP CDH1 CDH2 SNAI1 SNAI2 TWIST1
GSC
EMT
B
0Gy 2Gy 4Gy 8Gy 10Gy
Relative expression
PECAM1 CDH5 ENG VWF DES ACTA2 CD248 MCAM SOX2 NES
Endothelial
Pericyte
GSC
C
0Gy 8Gy_Day 2 8Gy_Day 3 8Gy_Day 7
% Double+ cells
CD31+ CD144+
CD146+ CD248+
D
CDH5-mC
DES-GFP
CDH5-mC DES-GFP
0 Gy
8 Gy
E
mC-
Des-mC+
CDH5-mC+
0 Gy
8 Gy
DESMIN
VE-CADHERIN
F
mC- CDH5-mc+ Des-mc+
Relative expression
CDH5 PECAM1 ENG DES ACTA2 PDGFRB SOX2
G
Pre-IR(0 Gy)
Post-IR(8 Gy)
DES-mc depleted
CDH5-mC depleted
H
Pre-depletion (0 Gy)
Post-depletion (0Gy_Day 3)
Post-depletion (8 Gy_Day 3)
%mC+ cells
DES-mC+
CDH5-mC+
I
0Gy_CDH5-mC+ 8Gy_CDH5-mC+
0Gy_DES-mC+ 8Gy_DES-mC+
%mC+ cells
HK_408 HK_157 HK_217 HK_417 HK_382 HK_372 HK_412
PN
CL
MES
J
CD133+
CD133-
* p<0.05, n=3
K
CD133+
CD133-
Log10 RLU
CD133+ CD133-
L
CD31 GFP
aSMA GFP
hGBM 0 Gy
hGBM 8 Gy
M
mGBM 0 Gy
mGBM 8 Gy
Relative expression
Pecam1 Cdh5 Eng Edn1 Des Mcam Cd248 Pdgfrb Sox2 Pou5f1 Nanog Klf4 Nestin Gfap
N
CD31 GFP
aSMA GFP
mGBM 0 Gy
mGBM 8 Gy

**Figure S2: Radiation promotes endothelial and pericyte marker expression**

(A) Graph shows relative expression of GSC and EMT genes in control (0 Gy), fractionated (2 Gy X4) and single dose (8 Gy) radiated glioma cells.

(B) Graph shows relative expression of endothelial, pericyte and GSC genes in control and low and high-dose radiated glioma cells.

(C) Flow cytometric quantitation of endothelial (CD31+ CD144+) and pericyte (CD146+ CD248+) markers in control and 2, 3 and 7 day-radiated glioma cells.

(D) Representative images of mCherry expression in control and radiated gliomaspheres. Scale bars, 150μm.

(E) Immunostaining VE-CADHERIN) and DESMIN in FACS sorted tumor (mC-) and transdifferentiated cells (CDH5-mC+/DES-mC+). Scale bars, 50μm.

(F) Quantitative RT-PCR analysis of endothelial, pericyte and GSC markers in sorted tumor (mC-) and transdifferentiated cells (CDH5-mC+ and DES-mC+) in control and radiated gliomaspheres.

(G,H) Images of mCherry in sorted, reporter-depleted control and radiated cells. Scale bars, 200μm. Graph shows quantitation of mCherry+ cells in each group.

(I) Flow cytometric quantitation of percentage of mCherry+ cells in endothelial and pericyte-reporter transduced control and radiated gliomaspheres from multiple patient-derived lines. PN-proneural, CL- classical and MES- Mesenchymal subtypes.

(J) Limiting dilution analysis of stem cell frequency in CD133+ and CD133- fractions

(K) Images of tumor growth in CD133+ and CD133- cells transplanted in mice. Graph shows quantitation of luminescence indicating tumor growth between the groups.

(L) Representative immunostaining images of CD31 and aSMA with GFP in tumor cells from control and radiated xenografts. Scale bars, 25um.

(M) Quantitative RT-PCR analysis of endothelial, pericyte and GSC markers in control and radiated mGBM cells. * and ** Indicates $p<0.05$, and $p<0.005$ in all graphs, two-tailed t-test.

#### Figure S3

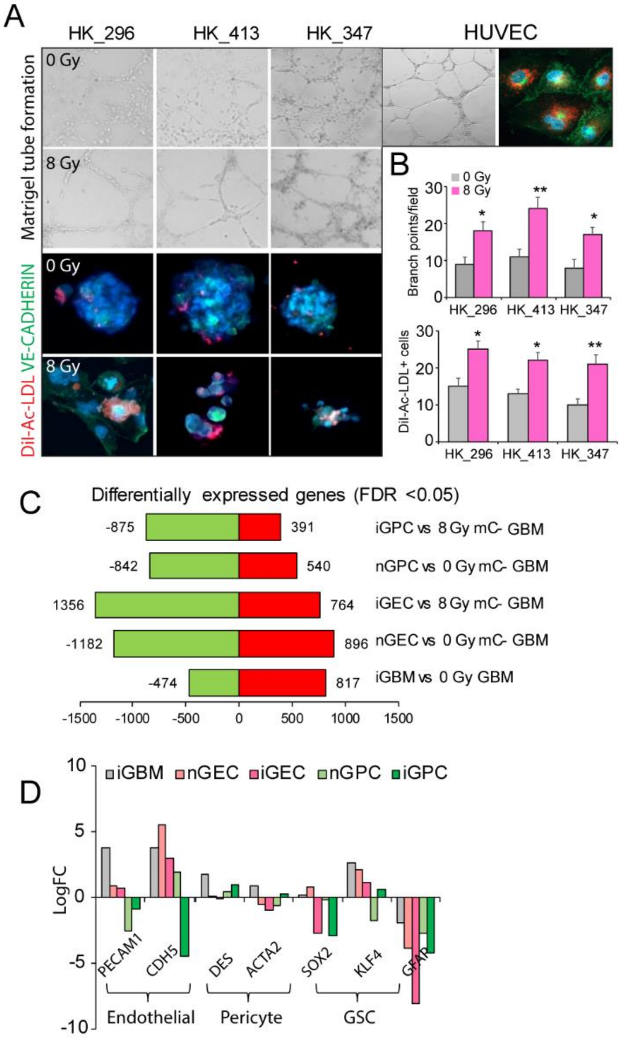


**Figure S3: iGEC show characteristics of normal vascular endothelial cells**

(A,B) Tubular network formation on matrigel and immunostaining of VE-CADHERIN and Di-AcLDL uptake of 7-day cultured control and radiated glioma cells from 3 patient-derived gliomaspheres lines. HUVEC were used as positive control. Scale bars, 100µm. Quantitation of branch points and DiI-Ac-LDL+ VE-CADHERIN+ cells per field is shown in the graph. * and ** Indicates p<0.05, and p<0.005, two-tailed t-test

(B) Graph shows the number of differentially expressed genes between GBM cells and transdifferentiated cells.

(C) Graph shows LogFC values of endothelial, pericyte and GSC genes expressed in the different fractions.

### Figure S4

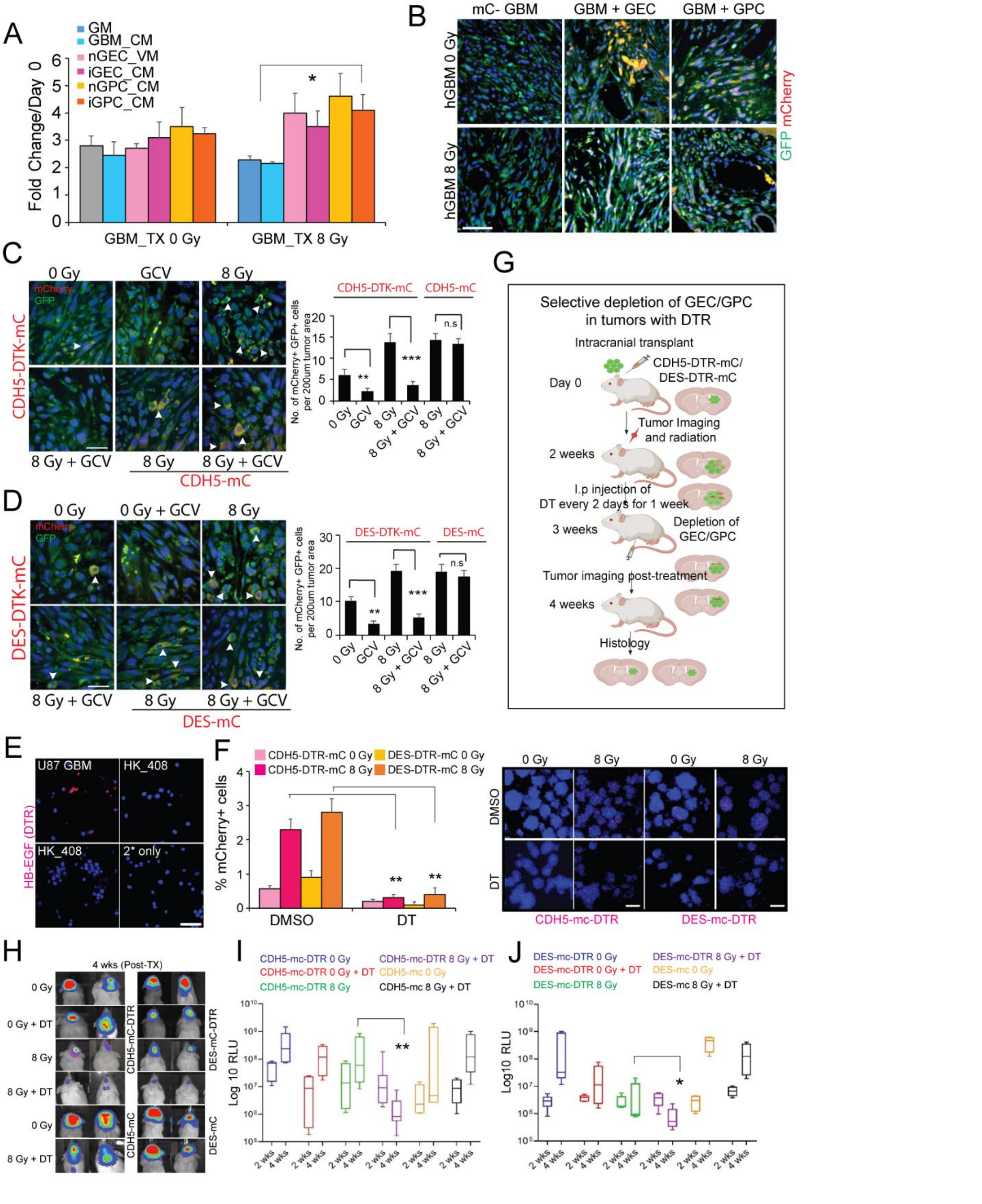


**Figure S4: Depletion of iGEC and iGPC inhibits tumor growth post-treatment**

(A) Graph shows quantitation of proliferation of sorted tumor cells from xenografts in GEC and GPC conditioned media. Error bars represent s.d. of mean and * $p<0.05$, two-tailed t-test.

(B) Immunostaining of GFP and mCherry in control and radiated tumors co-transplanted with GEC/GPC. Scale bars, 200um.

(C,D) Immunostaining of GFP and mCherry in control and radiated tumor xenografts treated with GCV to deplete GEC/GPC. Graphs show quantitation of number of GFP+ mCherry+ cells in each group. * and ** indicates p-value< 0.05 and <0.005, one-way ANOVA, post-hoc t-test

(E) Immunostaining of HB-EGF (DTR, red) in U87 GBM and HK_408 cells.

(F) Graph shows quantitation of mCherry+ cells pre-and post-radiation and diphtheria toxin (DT) treatment. Images show expression of mCherry in control and radiated cells after DMSO or DT treatment. Error bars represent mean ± SD, * indicates $p<0.05$, two-tailed t-test, N=3.

(G) Schematic outlines depletion of GEC/GPC using DTR-DT strategy in tumor xenografts

(H-J) Images of mice showing tumor growth post-treatment with DT. Box plots show the quantitation of tumor growth by luminescence pre- (2 week) and post-radiation and DT treatment (4 weeks). * indicates $p<0.05$ derived from one-way ANOVA and post-hoc t-test.

**Figure S5**

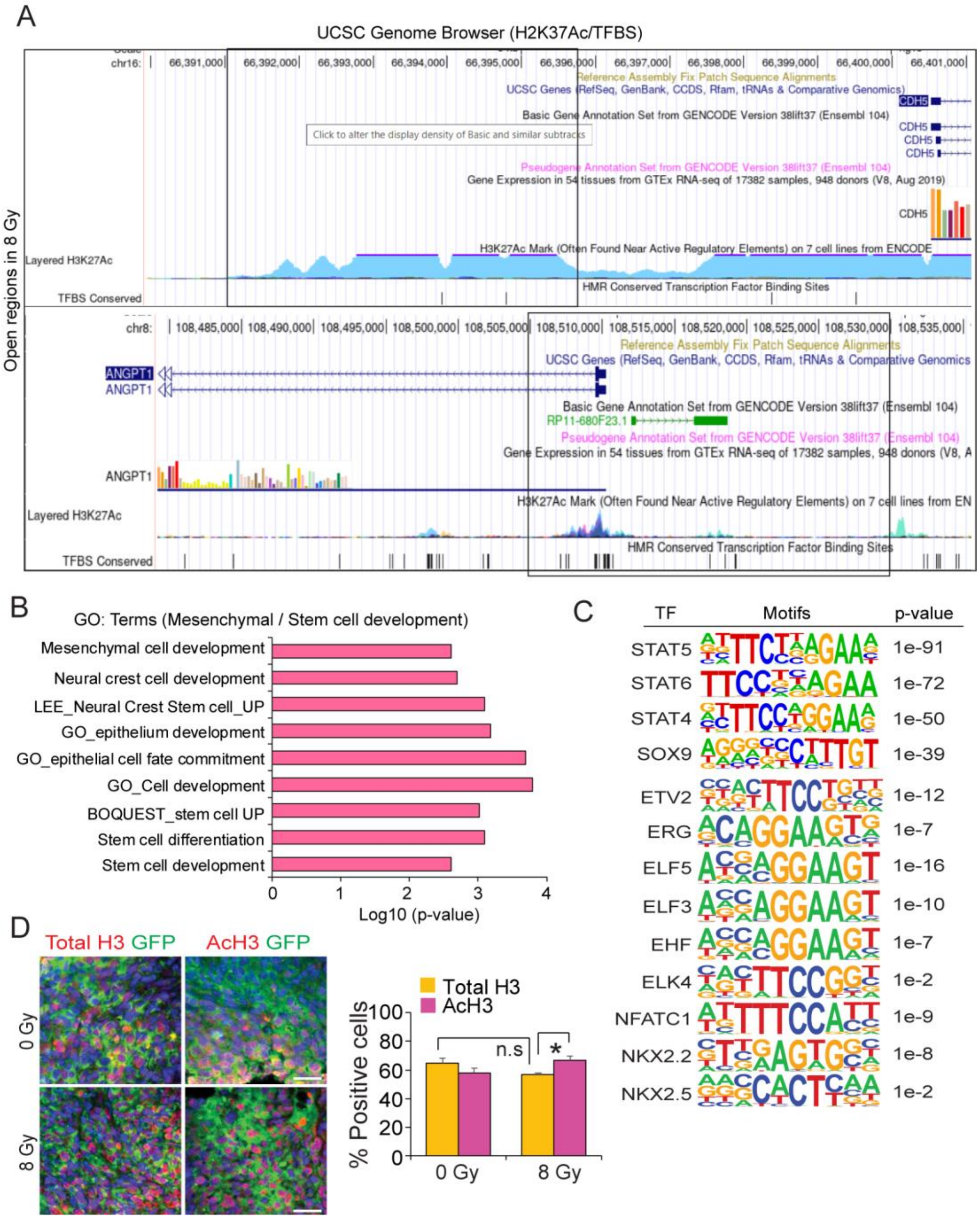


**Figure S5: Radiation alters chromatin accessibility in vascular gene regions**

(A) UCSC Genome Browser view of the CDH5 and ANGPT1 gene regions (hg19 assembly) with known H3K27Ac mark and TFBS sites. Region highlighted in black box is differentially open in radiated glioma cells.

(B) Graph shows significantly enriched gene ontology (GO) terms related to mesenchymal and stem cell development associated with genes differentially open in radiated gliomaspheres.

(C) Motifs significantly enriched in differentially open regions in radiated gliomaspheres. p-values are derived using hypergeometric or binomial distribution in HOMER.

(D) Immunostaining images of AcH3 and total H3 (red) in tumor cells (GFP, green) in control and radiated mouse GBM tumors. Quantitation of percentage of positive cells is shown in the graph. N=5 tumors per group, and * indicates $p<0.05$, two-tailed t-test. Scale bars, 100um.

**Figure S6**

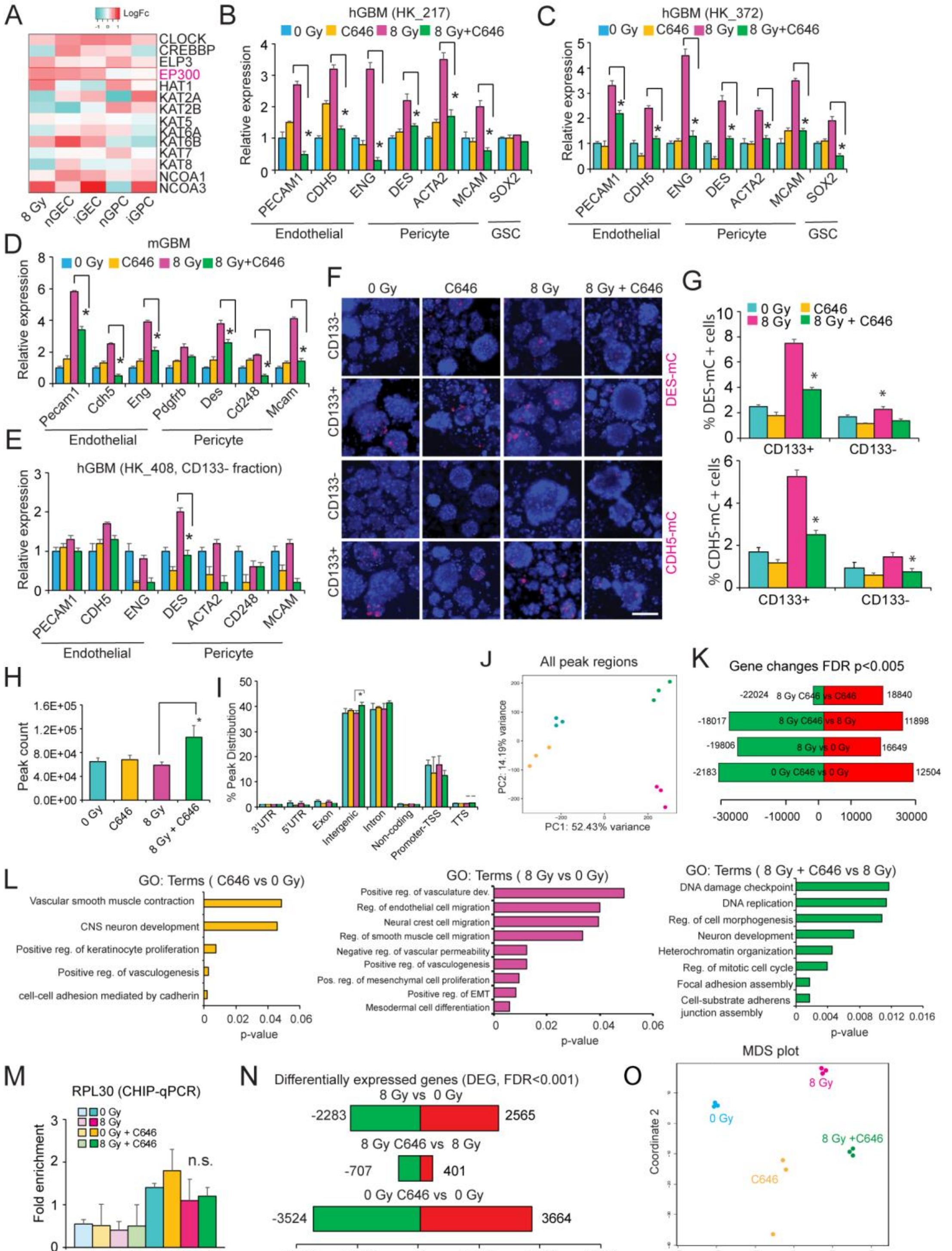
A
B
hGBM (HK_217)
C
hGBM (HK_372)
D
mGBM
E
hGBM (HK_408, CD133- fraction)
F
G
H
I
J
All peak regions
K
Gene changes FDR p<0.005
L
GO: Terms ( C646 vs 0 Gy)
GO: Terms ( 8 Gy vs 0 Gy)
GO: Terms ( 8 Gy + C646 vs 8 Gy)
M
RPL30 (CHIP-qPCR)
N
Differentially expressed genes (DEG, FDR<0.001)
O
MDS plot

**Figure S6: Blocking P300 HAT activity inhibits vascular marker expression**

(A) Heatmap shows expression of HAT transcripts from bulk RNA-sequencing of sorted tumor (8 Gy GBM) and transdifferentiated cells (nGEC/iGEC and nGPC/iGPC).

(B-E) Quantitative RT-PCR analysis of endothelial, pericyte marker expression in control and radiated hGBM lines (HK_217 and HK_372), mGBM and sorted CD133- hGBM fractions treated with C646. Error bars represent s.d and * indicates p<0.05 derived from two-tailed t-test.

(F,G) Images of mCherry+ expression in sorted CD133+ and CD133- fractions after radiation and C646 treatment. Graphs show quantitation of endothelial (CDH5-mC) and pericyte (DES-mC) reporter expression in each group.

(H, I) Average peak counts and peak distribution across various genomic regions from ATAC-sequencing of control (0 Gy), HAT inhibitor (C646), radiated (8 Gy) and combined radiation + HAT inhibitor (8 Gy+ C646) treated gliomaspheres. N=3 replicates per group, and * indicates p<0.05, derived from one-way ANOVA.

(J) PCA plot of all peak regions from control (0 Gy), HAT inhibitor (C646), radiated (8 Gy) and combined radiation + HAT inhibitor (8 Gy+ C646) treated gliomaspheres.

(K) Genes changes associated with peaks between control, radiated and C646 treated groups

(L) Gene ontology (GO) terms associated with differentially open regions in C646 treated and radiated cells.

(M) Fold enrichment of H3K27Ac in RPL30 gene in different groups. N=3 replicates and n.s. indicates not significant, two-tailed t-test.

(N) Graph shows differentially expressed genes between control, radiated and C646 treated groups from bulk RNA-sequencing.

(O) MDS plot of gene expression differences between control, radiated and C646 treated gliomaspheres.

**Figure S7**

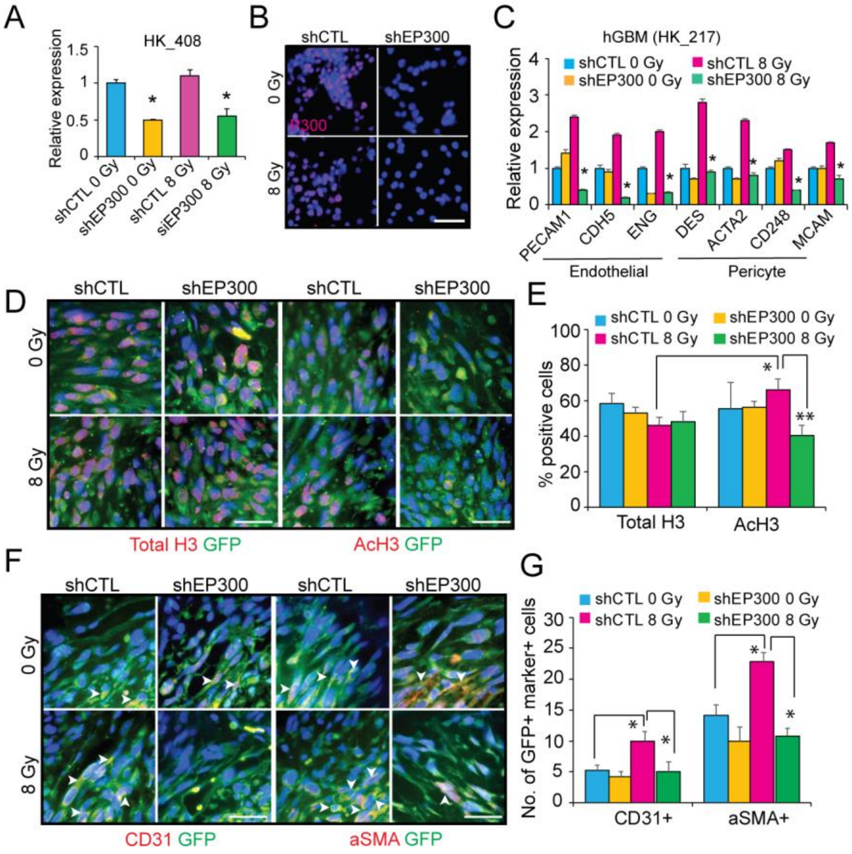


**Figure S7: EP300-deficient glioma cells show reduced H3K27Ac and vascular marker expression post-treatment**

(A) Quantitative RT-PCR analysis of EP300 transcript in control (shRNA-scrambled CTL) and knockdown (shRNA-EP300) cells. * indicates p<0.05, two-tailed t-test.

(B) Representative immunostaining images of P300 in control and knockdown cells.

(C) Quantitative RT-PCR analysis of endothelial and pericyte genes in control and knockdown glioma cells pre- and post-radiation. * indicates p<0.05, one-way ANOVA.

(D, E) Representative immunostaining images of Total H3 and AcH3, and GFP in control and knockdown tumor cells in control and radiated xenografts. Scale bars, 50μm Graph shows quantitation of marker+ GFP+ positive in each group. N=5 mice per group. * indicates p <0.05, one-way ANOVA.

(F,G) Representative immunostaining images of CD31 and aSMA and GFP in tumor sections. Scale bars, 50μm. Quantitation of GFP+ marker+ cells in tumor mass is shown in the graph. N=5 mice per group. * indicates $p < 0.05$, one-way ANOVA.
